# IER3IP1-mutations cause microcephaly by selective inhibition of ER-Golgi transport

**DOI:** 10.1101/2024.01.17.576044

**Authors:** Mihaela Anitei, Francesca Bruno, Christina Valkova, Therese Dau, Emilio Cirri, Iván Mestres Lascano, Federico Calegari, Christoph Kaether

**Author notes:** Correspondence should be addressed to: Christoph Kaether, Leibniz Institute on Aging, Fritz-Lipmann Institute, Beutenbergstr. 11, 07745 Jena, Germany.

## Abstract

Mutations in the *IER3IP1* (Immediate Early Response-3 Interacting Protein 1) gene can give rise to MEDS1 (Microcephaly with Simplified Gyral Pattern, Epilepsy, and Permanent Neonatal Diabetes Syndrome-1), a severe condition leading to early childhood mortality. The small endoplasmic reticulum (ER)-membrane protein IER3IP1 plays a non-essential role in ER-Golgi transport. Here, we employed secretome and cell-surface proteomics to demonstrate that the absence of IER3IP1 or the presence of the pathogenic p.L78P mutation results in the retention of specific cell-surface receptors and secreted proteins crucial for neuronal migration within the ER. This phenomenon correlates with the distension of ER membranes and increased lysosomal activity. Notably, the trafficking of cargo receptor ERGIC53 and KDEL-receptor 2 is compromised, with the latter leading to the anomalous secretion of ER-localized chaperones. Our investigation extended to in-utero knock-down of *Ier3ip1* in mouse embryo brains, revealing a morphological phenotype in newborn neurons. In summary, our findings provide insights into how the loss or mutation of a 10 kDa small ER-membrane protein can cause a fatal syndrome.

## Introduction

Microcephaly is characterized by a reduction of the head circumference by more than 2 standard deviations below the age and sex-corrected average. It can be caused by reduced proliferation or cell death of neuroprogenitor cells^1^, as well as by defective neuronal morphology or migration^2, 3^. Microcephaly, epilepsy, and diabetes syndrome-1 (MEDS1, OMIM #614231) is an autosomal recessive neurodevelopmental disorder characterized by microcephaly, simplified gyral pattern, severe epilepsy, and early-onset infantile diabetes mellitus caused by mutations in the *IER3IP1* gene^4^. Patients suffering from MEDS1 present dysmorphic facial features, skeletal deformities, and die in early childhood at the age of 1.5-8 years^5, 6^. Two point mutations in *IER3IP1* have been identified, p.V21G and p.L78P^4, 5, 7^. Additionally, one patient has been described with a compound heterozygosity in *IER3IP1*, with one allele carrying the p.V21G variant, and the other a c.79delT deletion (p.T79Δ) resulting in a frameshift mutation (p.Phe27fsSer*25)^6^. Recently, a MEDS1 patient with a third, homozygous, p.A18V variant has been detected. IER3IP1 is an 82-amino acid protein with two predicted transmembrane domains, that localizes to the endoplasmic reticulum (ER)^4,9^. In mouse, Ier3ip1 is highly expressed during embryogenesis, especially at sites of neurogenesis (i.e., cortical ventricular and subventricular zones)^4^. Its yeast orthologue, Yos1p, together with Yip1p and Yif1p, forms a tripartite complex essential for ER-Golgi transport^10^. Interestingly, mutations in the human orthologue of *Yip1p*, *YIPF5*, cause MEDS2 (OMIM #614231), a syndrome similar to MEDS1^11^. The third complex subunit, *Yif1p*, has two human orthologues, *YIF1A* and *YIF1B*, and *YIF1B* mutations can also cause microcephaly and epilepsy^12^, suggesting common pathomechanisms of the *YIPF5*, *YIF1B* and *IER3IP1* disease-causing variants.

At the cellular level, loss of IER3IP1 induces apoptotic cell death in pancreatic β-cells^4^ and in mouse insulinoma cells^13^, where it also impairs the reaction to ER stress, the unfolded protein response (UPR). A mouse model with β-cell-specific deletion of Ier3ip1 presents, in contrast, increased ER-stress without an evident UPR activation^14^. The ER is dilated in these cells, and proinsulin folding is impaired. In human brain organoids, IER3IP1 determines size and extracellular matrix composition^15^. These studies indicate an important role for IER3IP1 in the ER, however, its function in ER to Golgi transport and its potential cargos in mammalian cells are not well understood. We here provide a detailed analysis of IER3IP1 function and show that it governs the ER-export of a subset of secretory and membrane-bound cargos. The ER-export of several factors important for neuronal migration and axon pathfinding is reduced, and the localization of ERGIC53 and KDEL-receptor 2 are changed, the latter causing an abnormal secretion of ER-localized enzymes. We also provide a comparative functional study of known pathogenic variants, which indicate that *IER3IP1* p.L78P, p.T79Δ, but not p.V21G, disrupt the ER-export of specific cargos. Finally, we model the loss of IER3IP1 during mammalian brain development by in utero electroporation showing its importance for coordinating neuronal morphology.

## Results

To investigate the specific role of IER3IP1 in the early secretory pathway, we used CRISPR/Cas9 to knock out (KO) this protein in HeLa cells (Fig. 1A). Two monoclonal cell lines (KO1 and KO2) were selected. The *IER3IP1* gene was fully deleted in the KO1 clone (Fig. 1B-E), and indels were present in the KO2 cells (Fig. 1B, C). A control clonal cell line was established from a pool of cells transfected with Cas9 only. IER3IP1 protein loss was confirmed by immunoblotting (Fig. 1K), using a specific antibody against IER3IP1. Cell growth of both KO1 and KO2 cells, determined by Incucyte live-cell imaging for 72 h, was not significantly different compared to controls (Fig. 1F), suggesting IER3IP1 was not essential for HeLa cell survival. Since patient-derived cells were not accessible, we re-expressed wild-type (WT), p.L78P, p.V21G or p.T79Δ mutants (schematized in Fig. 1J) in KO1 cells, and clonal cell lines for each variant were selected for further analysis. Re-expressed *IER3IP1* WT, p.L78P and p.V21G proteins were detectable by immunoblot (Fig. 1K) and by immunofluorescence analysis (Fig. 1G, H). In contrast, p.T79Δ protein was not detectable at steady state (Fig. 1K), or after incubation with a proteasome (MG132) or a lysosome (chloroquine) inhibitor (Suppl. Fig. S1A), although p.T79Δ *IER3IP1* mRNA was present in these cells (Suppl. Fig. S1B). Together, these data suggest that expression of the *IER3IP1* p.T79Δ variant is equivalent to a KO of *IER3IP1*.

**Figure 1.**
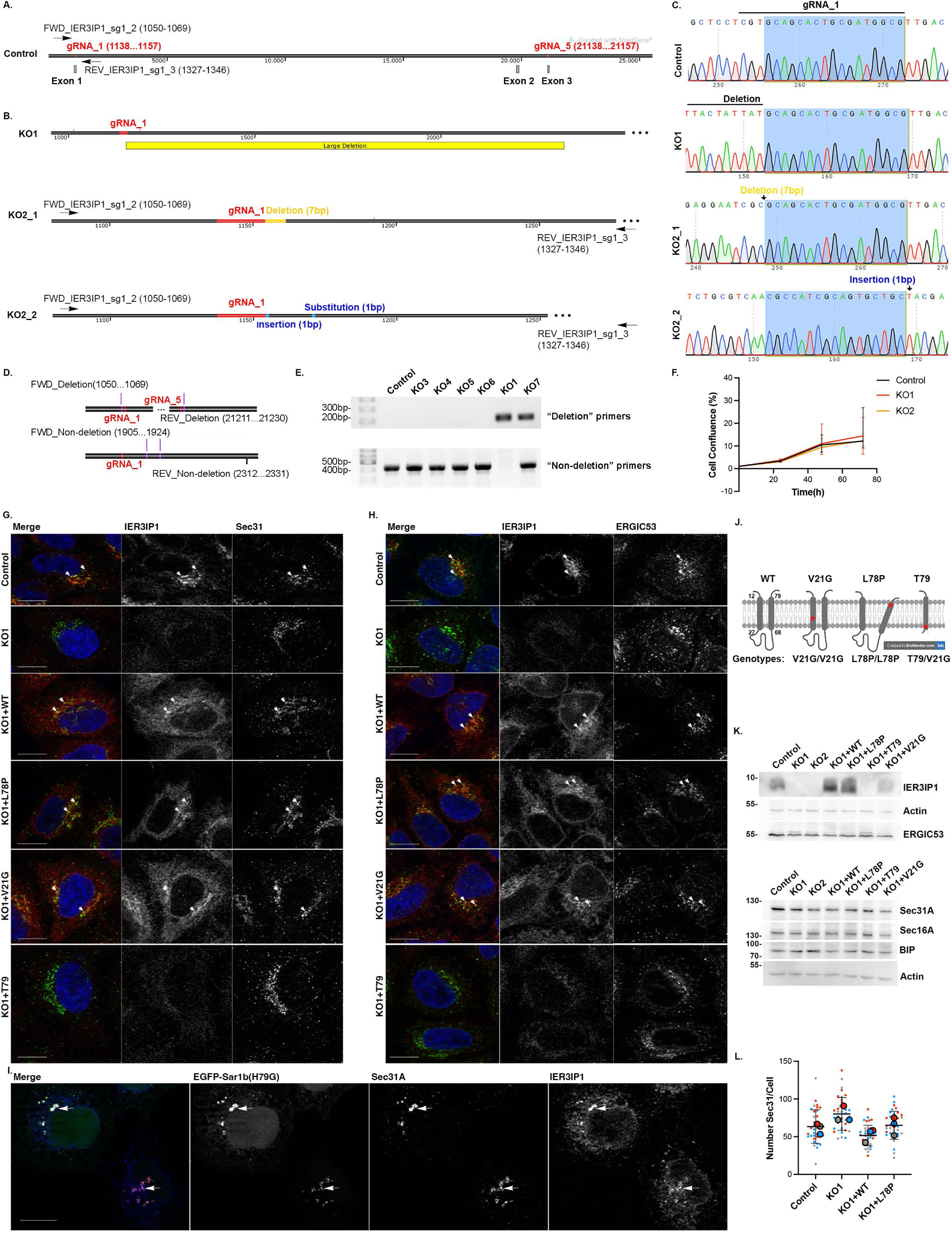
Characterization of CRISPR/Cas9-generated *IER3IP1* KO and mutant re-expressing HeLa cell lines. **(A)** Two guide RNAs (gRNA_1, gRNA_5) and the primers used to amplify the gRNA_1 cutting site for further characterization are shown. Models indicate **(B, C)** heterozygous indels in the KO2 clone, the large deletion in the KO1 clone, and **(D, E)** the localization of the “Deletion” and “Non-deletion” primers (red = base pairs targeted by gRNA_1). **(F)** Incucyte growth curve analyses of the indicated cell lines. Data were normalized to t0_Control_ and are shown as median±range (n = 4 independent experiments). **(G, H)** Cells were fixed, co-labeled with anti-IER3IP1 (red) and **(G)** Sec31 (green) or **(H)** ERGIC53 (green), and Hoechst nuclear dye (blue), and analyzed by fluorescence microscopy. Arrows indicate overlap of IER3IP1 with the respective marker. **(I)** Cells were transfected with EGFP-Sar1b (H79G), fixed after 24 h, labeled with anti-IER3IP1 (blue) and Sec31A (red), and analyzed by fluorescence microscopy. Arrows indicate co-localization of the three proteins. **(G-I)** Scale bars, 10µm. Single Apotome sections are shown. **(J)** Model of the IER3IP1 potential transmembrane domains and the localization of the pathogenic mutations. **(K)** *IER3IP1* KO and re-expression of *IER3IP1* variants in the indicated cell lines were analyzed by WB using anti-IER3IP1. Lysates were assessed for expression of the indicated proteins, with β-actin serving as loading control (n = 3 independent experiments). **(L)** The numbers of Sec31 objects per cell in cells processed as in (G) were analyzed using Cell Profiler. Values obtained from individual images are represented as small circles and mean values for independent experiments as filled circles (n = 3 independent experiments, mean ± SD, One-way Anova with Dunnett’s post-hoc test, n _Control_ = 506, n _KO1_ = 534, n _KO1+WT_ = 644, n _KO1+P.L78P_ = 493 cells).

Endogenous IER3IP1 was particularly enriched in the perinuclear region, where it partially overlapped with ER exit-sites (ERES) labeled by the COPII components Sec31A (Fig. 1G) or mCherry-Sec24C (Suppl Fig. S1D, E). Partial colocalization was also detected with ERGIC53, a marker of the ER-Golgi intermediate compartment (Fig. 1H), EGFP-ERGIC53 (Suppl Fig. S1D, E) and ER marked by mRFP-KDEL (Suppl Fig. S1C). An EGFP-tagged GTP-locked Sar1a (H79G) mutant, which arrests Sec31A on ERES and blocks ER-export^16^, increased colocalization of Sec31A with IER3IP1 (Fig. 1I), indicating that IER3IP1 transits through COPII-coated membranes. Exogenous IER3IP1 WT, p.L78P and p.V21G were also detectable in the perinuclear region, and showed an ER-like reticular pattern, similar to the endogenous protein (Fig. 1G, H). In conclusion, IER3IP1 is a component of the early secretory pathway that cycles between ER, ERGIC and the Golgi apparatus. IER3IP1 is dispensable for cell growth or survival, and MEDS1-causing mutations do not cause obvious changes in its localization.

We further asked if *IER3IP1* deletion affected the organization of the early secretory pathway. The distribution pattern and the number of Sec31A-labelled ERES^17^ in *IER3IP1* KO1 or mutant-expressing cells were similar to those in control or *IER3IP1* WT expressing cells (Fig. 1G, L). In contrast, the number of peripheral ERGIC53, but not Sec16A (another ERES marker)^17^ (Fig. 2A-C), puncta/cell was higher in *IER3IP1* KO1 cells, a change rescued by re-expression of the WT and p.V21G, but not of the p.L78P and p.T79Δ variants. Moreover, IER3IP1 WT, but not p.L78P, co-immunoprecipitated with EGFP-ERGIC53 from cell lysates (Fig. 2D), suggesting that IER3IP1 binds to ERGIC53 and participates in its exit from the ER.

**Figure 2.**
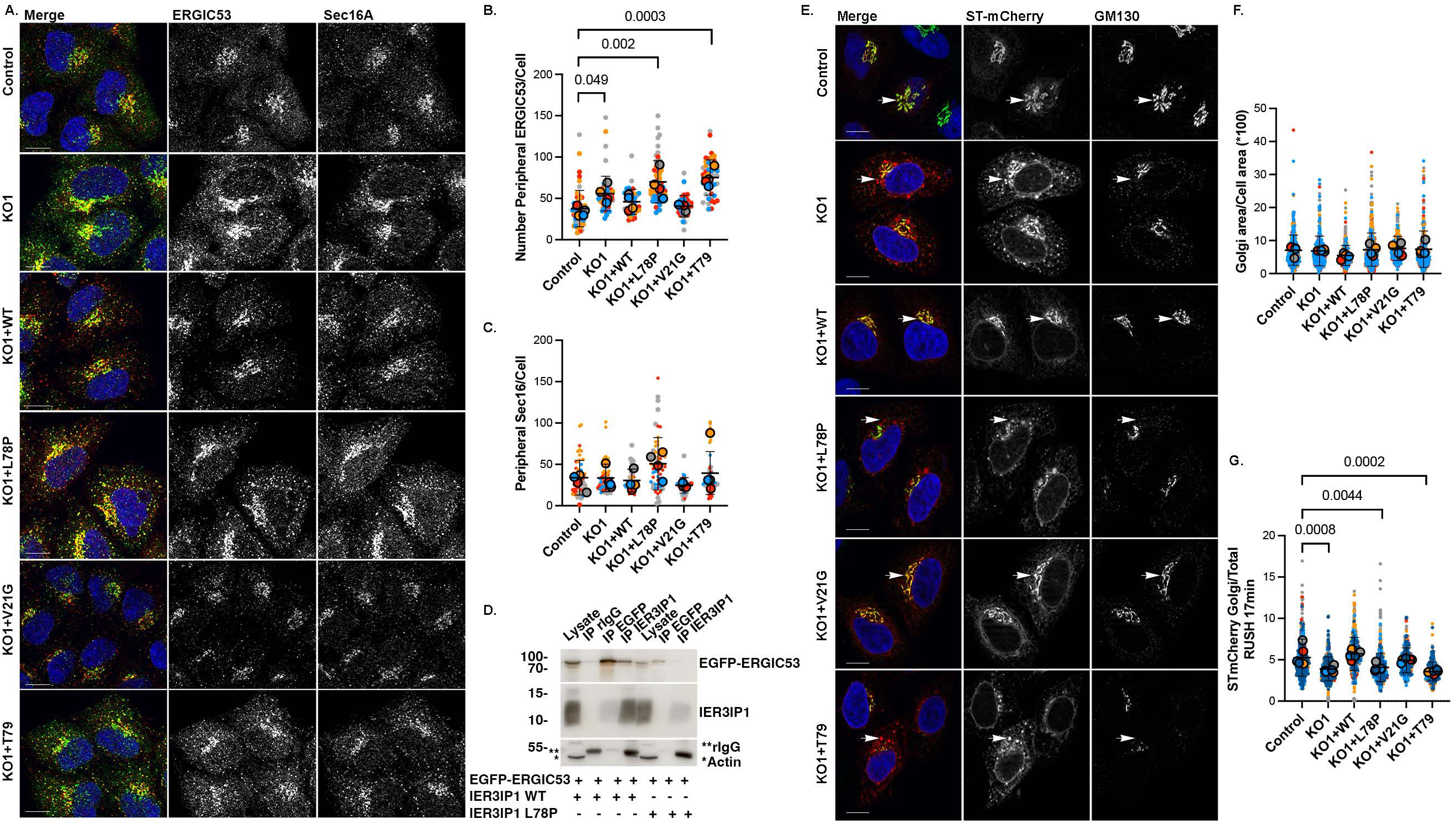
Cargo exit from the ER is delayed in *IER3IP1* KO cells and in cells expressing the *IER3IP1* p.L78P and p.T791′ pathogenic mutations. **(A-C)** Cells fixed and co-labeled with anti-ERGIC53 (green), anti-Sec16A (red) and Hoechst (blue) were analyzed by fluorescence microscopy. **(B, C)** ERGIC53 **(B)** or Sec16A **(C)** objects were counted in the cell periphery, excluding the perinuclear subpopulation, using Cell Profiler. Measurements from individual images (small circles), mean values per experiment (large, filled circles) are indicated, with biological replicates color-coded (n = 4, except for n _KO1+p.V21G_ = 3, mean ± SD, One-way Anova with Dunnett’s post-hoc test; n _Control_ = 896, n _KO1_ = 831, n _KO1+WT_ = 566, n _KO1+p.L78P_ = 766, n _KO1+p.V21G_ = 513, n _KO1+p.T79Δ_ = 614 cells). **(D)** HeLa cells were transfected with the indicated constructs. IER3IP1 and EGFP-ERGIC53 were immunoprecipitated from the lysates using anti-IER3IP1 or anti-EGFP antibodies, and then analyzed by immunoblotting (n = 5 independent experiments). **(E-G)** Cell lines were transfected with Str-STIM-ST-SBP-mCherry (ST-mCherry). After 24 h, cells were incubated with biotin for 17 min, fixed and co-stained with anti-GM130 (Golgi marker, green) and Hoechst dye (blue), and analyzed by fluorescence microscopy. **(F)** The areas of GM130-stained Golgi measured with Fiji in individual cells (small circles), and the color-coded mean values per experiment (large, filled circles) are shown (n = 4 independent experiments, mean ± SD, One-way Anova with Dunnett’s post-hoc test; n _Control_ = 476, n _KO1_ = 435, n _KO1+WT_ = 418, n _KO1+p.L78P_ = 550, n _KO1+p.V21G_ = 252, n _KO1+p.T79Δ_ = 312 cells). **(G)** Ratios between Golgi and total (entire cell) ST-mCherry mean fluorescence intensities, measured using Fiji, are shown for individual cells (small circles), and for independent experiments (mean values indicated as large, filled circles) (n = 5 independent experiments, mean ± SD, One-way Anova with Dunnett’s post-hoc test; n _Control_ = 515, n _KO1_ = 475, n _KO1+WT_ = 476, n _KO1+p.L78P_ = 624, n _KO1+p.V21G_ = 293, n _KO1+p.T79Δ_ = 368 cells). **(A, E)** Scale bars, 10µm. Single Apotome sections are shown.

The steady state morphology of cis-Golgi, labeled with an antibody against GM130, was not affected in IER3IP KO or mutant-expressing cells (Fig. 2E, F). To test if IER3IP1 played a role in *de novo* formation of Golgi stacks, cells were incubated for 1 h with brefeldin A (BFA), a compound that leads to fusion of Golgi membranes with the ER^18^. BFA washout allowed Golgi reassembly, as observed after 2 h (Suppl. Fig. S2A-D). Compared to controls, the formation of novel Golgi stacks was significantly reduced in *IER3IP1* KO1 cells, and it was rescued by re-expression of the WT or p.V21G, but not of the p.L78P or p.T79Δ IER3IP1 variants (Suppl. Fig. S2D). Moreover, after Golgi reassembly, the constitutive Golgi transmembrane protein ST6GAL1 co-localized less with GM130 in the absence of IER3IP1 (Suppl. Fig. S2E), suggesting ST6GAL1 was transported slower from the ER to Golgi than in control cells. This delay was rescued by WT and p.V21G, but not by p.L78P or p.T791′ IER3IP1. To confirm that IER3IP1 was important for ST6GAL1 transport, we used the retention using selective hook (RUSH) assay^19^. Cells were transfected with a plasmid (ST-mCherry) encoding ST6GAL1 fused to streptavidin binding peptide and mCherry, and a hook composed of streptavidin fused to an ER retention signal (Str-STIM1)^19^. Upon biotin addition, the interaction between SBP and streptavidin was released, allowing the transport of ST-mCherry from the ER to the Golgi. 17 min after biotin addition, the ratio between Golgi-localized and total cellular ST-mCherry was reduced in IER3IP1 KO1 and KO2 compared to control cells, suggesting a delay in the synchronized transport of ST-mCherry from the ER to the Golgi (Fig. 2E, G). This reduced transport was rescued by re-expression of WT and p.V21G, but not by p.L78P and p.T79Δ. In sum, IER3IP1 is involved in ER-Golgi transport of ERGIC53 and the Golgi membrane protein ST6GAL1, and the p.L78P and p.T79Δ variants impair this function.

### IER3IP1 controls ER export of specific plasma membrane proteins

To get more insights into the role of IER3IP1 in the transport of endogenous cargo, we isolated the surface proteomes of control, *IER3IP1* KO1, and WT *IER3IP1* re-expressing HeLa cells, utilizing cell surface biotinylation and mass spectrometry (MS). About 3900 proteins were detected for each condition, with a high reproducibility (Suppl. Fig. S1O). In biotinylated control cells, 1006 proteins were significantly enriched over non-biotinylated controls, 31% of which could be GO term-annotated as components of either the plasma membrane, cell surface, extracellular space, extracellular matrix or extracellular exosome, 31% as transmembrane proteins, and 35% as glycoproteins (Fig. 3A, Suppl. Table S1). A total of 235 surface proteins were differentially expressed in *IER3IP1* KO1 cells compared to controls (Suppl. Table S2). Among these, 90% were not changed in the total proteome of *IER3IP1* KO1 ve*rsus* control cells (Suppl. Tables S2, S3), suggesting only their transport was affected (Fig. 3B). Re-expression of *IER3IP1* WT rescued the surface levels of most proteins (Fig. 3C). Pathway enrichment analysis revealed that the reduced proteins on the surface of *IER3IP1* KO1 cells compared to controls (Fig. 3D-G, Suppl. Table S2) were linked to the immune system and to nervous system development. Various proteins implicated in neuronal function were either increased, i.e. integrins (ITGA3, ITGA5, ITGB1), the semaphorin receptor neuropilin 1, or decreased, i.e. FGFR3 and FGFR2, the netrin 1 receptor UNC5B, the hepatocyte growth factor receptor MET, EPHB3 and laminin subunit alpha 1 (LAMA1). Among the proteins that could be associated with either apical or basolateral plasma membrane domains (http://polarprotdb.ttk.hu/), we observed an enrichment (76.4%) in basolateral proteins on the surface of *IER3IP1* KO1 cells compared to control (Suppl. Table S7). Total proteome analysis showed that cells lacking IER3IP1 downregulated components of the endo-lysosomal compartment, and upregulated membrane trafficking components, including the COPI subunits COPG1 and COPB2, the p24 protein TMED7 and small GTPases such as RAB6A and RAB11B (Suppl. Table S3, Suppl. Fig. 1E-I), maybe to compensate the absence of IER3IP1.

**Figure 3.**
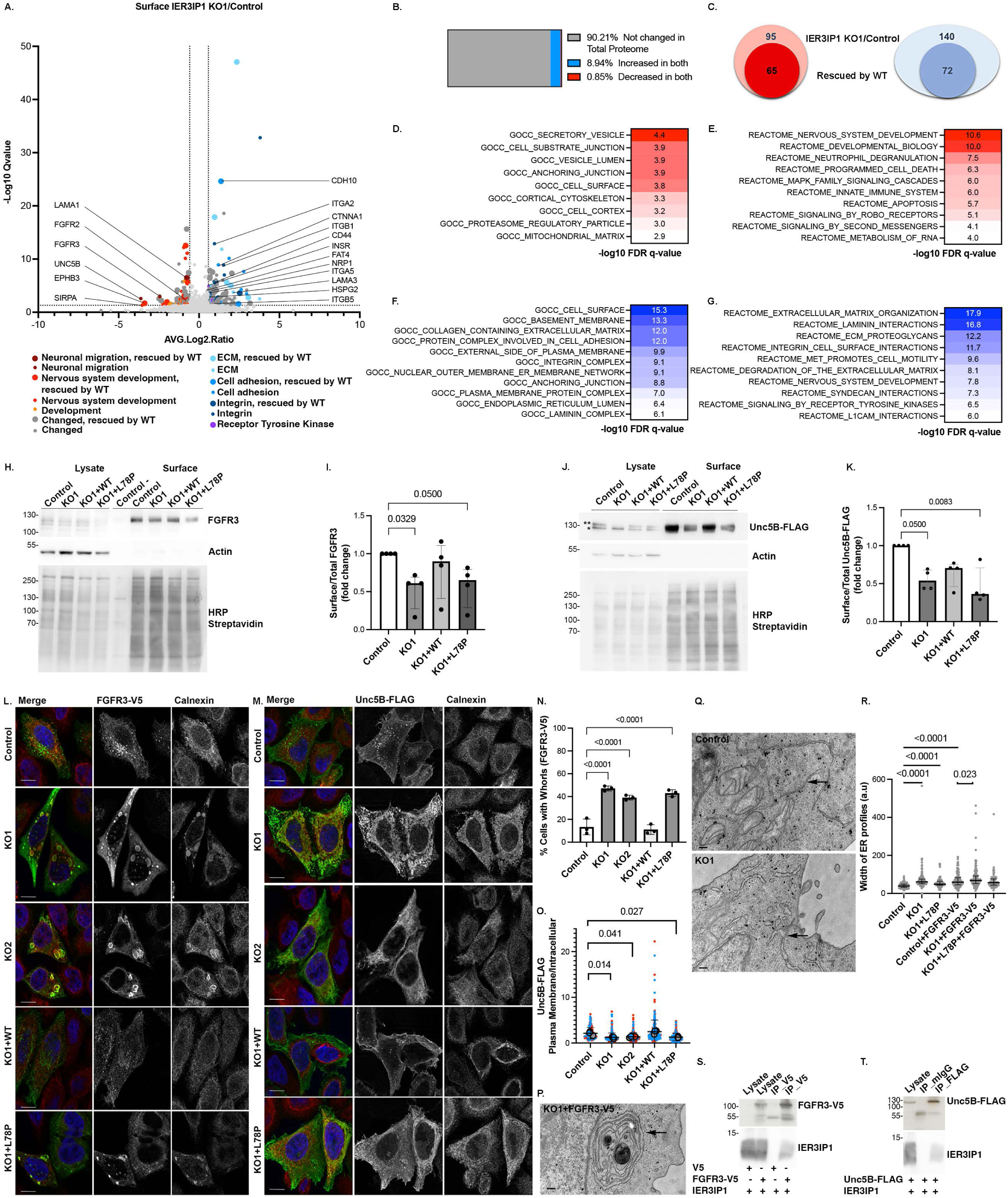
*IER3IP1* KO changes the composition of the surface proteome. **(A)** Volcano plot of proteins differentially regulated on the surface of *IER3IP1* KO1 and control HeLa cells (Suppl. Table S2). Association with various biological pathways was color-coded (legend). Large circles indicate proteins rescued by *IER3IP1* WT (n = 3 independent experiments). **(B)** Comparison of proteins modified on cell surface and in the total proteome (Suppl. Tables S2, S3). **(C)** Number of proteins decreased (red) or increased (blue) on the surface of *IER3IP1* KO1 cells compared to controls (outer circle) and rescued by *IER3IP1* WT (inner circle). **(D-G)** GSEA pathway enrichment analysis of hits either reduced (red) or increased (blue) in KO1 *vs* control (Suppl. Table S2). **(H-K)** Immunoblot analyses of endogenous FGFR3 **(H, I)** and Unc5B-FLAG **(J, K)**. Proteins were detected with anti-FGFR3, anti-FLAG, biotinylated proteins with HRP-conjugated streptavidin, and beta-actin was used as loading control. N = 4 independent experiments; median ± IQR, Kruskal-Wallis with Dunn’s post-hoc test. **(L-O)** Cells were transfected with **(L)** FGFR3-V5 or **(M)** Unc5B-FLAG, fixed and labeled with **(L)** anti-V5 or **(M)** anti-FLAG (green), anti-calnexin (red) and Hoechst (blue). Scale bars, 10 µm. Single Apotome sections are shown. **(N)** Percentage of cells expressing FGFR3-V5 and displaying whorls (n = 3 independent experiments; mean ± SD, n ≥ 351 cells/condition). **(O)** Surface/intracellular Unc5B-FLAG fluorescence intensity ratios are shown per cell (circles), and per experiment (filled circles) (n = 3, mean ± SD; n ≥ 173 cells/condition). Statistical significance was analyzed using One-way Anova with Dunnett’s post-hoc test. **(P-R)** Control and *IER3IP1* KO1 cells were fixed and processed for transmission electron microscopy. Arrows indicate a whorl **(P)** or ER profiles **(Q)**. Scale bars, 200 nm. **(R)** Width of ER profiles was analyzed with Fiji (n ≥ 10 cells/condition; n ≥ 74 ER profiles; median ± IQR; Kruskal-Wallis with Dunn’s post-hoc test). **(S, T)** Transiently expressed FGFR3-V5 **(S)** and Unc5B-FLAG **(T)** were immunoprecipitated from lysates using anti-V5 and anti-FLAG, respectively, and analyzed by immunoblotting (n = 6 independent experiments).

Subsequent validation by immunoblotting confirmed that surface levels of endogenous FGFR3 and exogenous Unc5B-FLAG cells were reduced in *IER3IP1* KO1 compared to controls (Fig. 3H-K, Suppl. Fig. S1F, G). Interestingly, these changes were partially rescued by re-expressing *IER3IP1* WT, but not the pathogenic mutant p.L78P. Moreover, mature, complex glycosylated Unc5B-FLAG, indicated by the slower migrating band and representing the surface-localized protein, was significantly reduced in KO1 cells compared to controls. Consequently, the proportion of the lower, immature band (representing the ER-localized Unc5B-FLAG), relative to the total protein amount, was higher. Changes in Unc5B-FLAG glycosylation were rescued by IER3IP1 WT (Suppl. Fig. S1H, I). These data concur with immunofluorescence microscopy analysis showing that the receptor was retained in the ER (identified with an antibody against the ER marker calnexin) in *IER3IP1* KO1 and KO2 cells, whereas it was mostly detected on the cell surface in control and *IER3IP1* WT cells (Fig. 3M, O). *IERIP1* p.L78P did not rescue the transport of Unc5B-FLAG to the plasma membrane (Fig. 3M, O), although it did restore its glycosylation (Suppl. Fig. S1H, I), suggesting a significant amount of receptor was transported through the Golgi apparatus in these cells. FGFR3-V5 localization was also affected in *IER3IP1* KO1, KO2 and p.L78P cells, where the receptor was detectable in ER membrane “whorls” co-labeled with the ER-marker calnexin (Fig. 3L, N). FGFR3-V5 and Unc5B-FLAG pulled down co-expressed IER3IP1 from cell lysates (Fig. 3S, T), suggesting that IER3IP1 interacts with both and controls their ER export.

Using transmission electron microscopy, we confirmed that ER whorls were only detectable in FGFR3-V5 expressing KO1 and p.L78P cells (Fig. 3P, Q). Independently of FGFR3-V5 expression, ER stacks appeared enlarged in *IER3IP1* KO1 and p.L78P cells *versus* controls (Fig. 3Q, R). Changes in ER shape were further evaluated by live fluorescence microscopy analysis of cells overexpressing the ER membrane protein P180 (Ribosome Binding Protein 1, RRBP1). EGFP-p180-labeled ER membranes covered a significantly higher percentage of the cell area in *IER3IP1* KO1 and p.L78P compared to controls and WT expressing cells (Suppl. Fig. S3A, B). This suggests that, in the absence of IER3IP1, increased protein accumulation in the ER causes membranes to enlarge and, in some cases, to form whorls to accommodate cargo excess.

Whorl formation may represent a novel type of ER stress response that inhibits protein translation by disentangling translocons from ribosomes^20^. Immunoblot analysis of cells overexpressing FGFR3-V5, however, did not show any changes in the levels of the ER stress sensor BiP in the absence of IER3IP1 (Suppl. Fig. S3C, D). Moreover, BiP upregulation was similar in all tested cell lines after incubation with thapsigargin, an ER stress inducer (Suppl. Fig. S3 E, F). The levels of phospho-IRE1 (pIRE-1) (Suppl. Fig. S3G) and ATF6 (Suppl. Fig. S3H, I), proteins involved in the UPR response, were not changed either, compared to controls, at steady state or after incubation with thapsigargin. These findings were consistent with the mass spectrometry analysis of the total proteomes, where no significant upregulation of the ER stress response was observed in cells lacking IER3IP1 (Suppl. Table S3). In summary, IER3IP1 may affect the homeostasis of the endo-lysosomal compartment and it selectively controls the surface transport of a subset of membrane proteins, including FGFR3 and UNC5B. Does IER3IP1 also control the secretion of soluble cargos?

### IER3IP1 controls the secretion of specific cargos

To find proteins whose secretion may depend on IER3IP1, we used click-chemistry to isolate biotinylated, metabolically labeled, newly synthesized glycoproteins from concentrated cell culture medium^21^. A total of 816 proteins were identified by MS analysis, with a high reproducibility (Suppl. Fig. S1P). Among these, 387 proteins were significantly enriched in the biotinylated over the non-biotinylated controls (Suppl. Table S4). Of these, 78.3% were glycoproteins, 80% could be GO term-annotated as either plasma membrane, cell surface, extracellular space, extracellular matrix or extracellular exosome, 36.17% as secreted, 39.27% as transmembrane proteins, and 69.25% contained a signal peptide bearing information for secretion (Suppl. Table S4). Overall, 221 proteins were differentially secreted by *IER3IP1* KO1 cells when compared to controls (Fig. 4A, Suppl. Table S5), most of which (95%) were not changed in the total proteome (Fig. 4B, Suppl. Tables S3, S5). Interestingly, *IER3IP1* KO reduced the secretion of SERPINA1 (alpha-1-antitrypsin), an established ERGIC53 cargo^22^, strengthening the hypothesis that IER3IP1 and ERGIC53 cooperate during membrane transport (see Fig. 2). Gene set enrichment analysis (GSEA) showed that *IER3IP1* deletion reduced the secretion of proteins belonging to pathways such as innate immune system, developmental biology, or intercellular junctions (Fig. 3D-G, Suppl. Table S5). Significantly, these included factors linked to neuronal migration, such as SERPING1, PAICS, CELSR1, LAMA1, SPARCL1 and APP. Proteins whose secretion was increased in the absence of IER3IP1 belonged to pathways like ECM organization (Fig. 4F, G), in agreement to the surfaceome results (Fig. 3F, G). Interestingly, several lysosomal enzymes were abnormally secreted by KO1 cells (Fig. 3A, G; Suppl. Table S5). To check the integrity of the lysosomal compartment in these cells, we used high content confocal fluorescence microscopy (Suppl. Fig. S4). Although the total numbers of LAMP1-positive late endosome/lysosomes were similar (Suppl. Fig. S4B, D), *IER3IP1* KO1 and p.L78P cells had significantly more active lysosomes labeled by Magic Red Cathepsin L than controls (Suppl. Fig. S4A, C). This increase in lysosome activity may be necessary for the clearance of excessive ER membranes^23^ observed in cells lacking functional IER3IP1. Assessed by immunoblotting, the amounts of the autophagosomal proteins LC3B and p62/SQSTM1^24^ were similar to controls in all tested experimental conditions (steady state, Torin1-dependent autophagy induction, chloroquine (CQ)-inhibition of lysosomal function (Fig. S4E-G), suggesting autophagy was not affected in the absence of IER3IP1.

**Figure 4.**
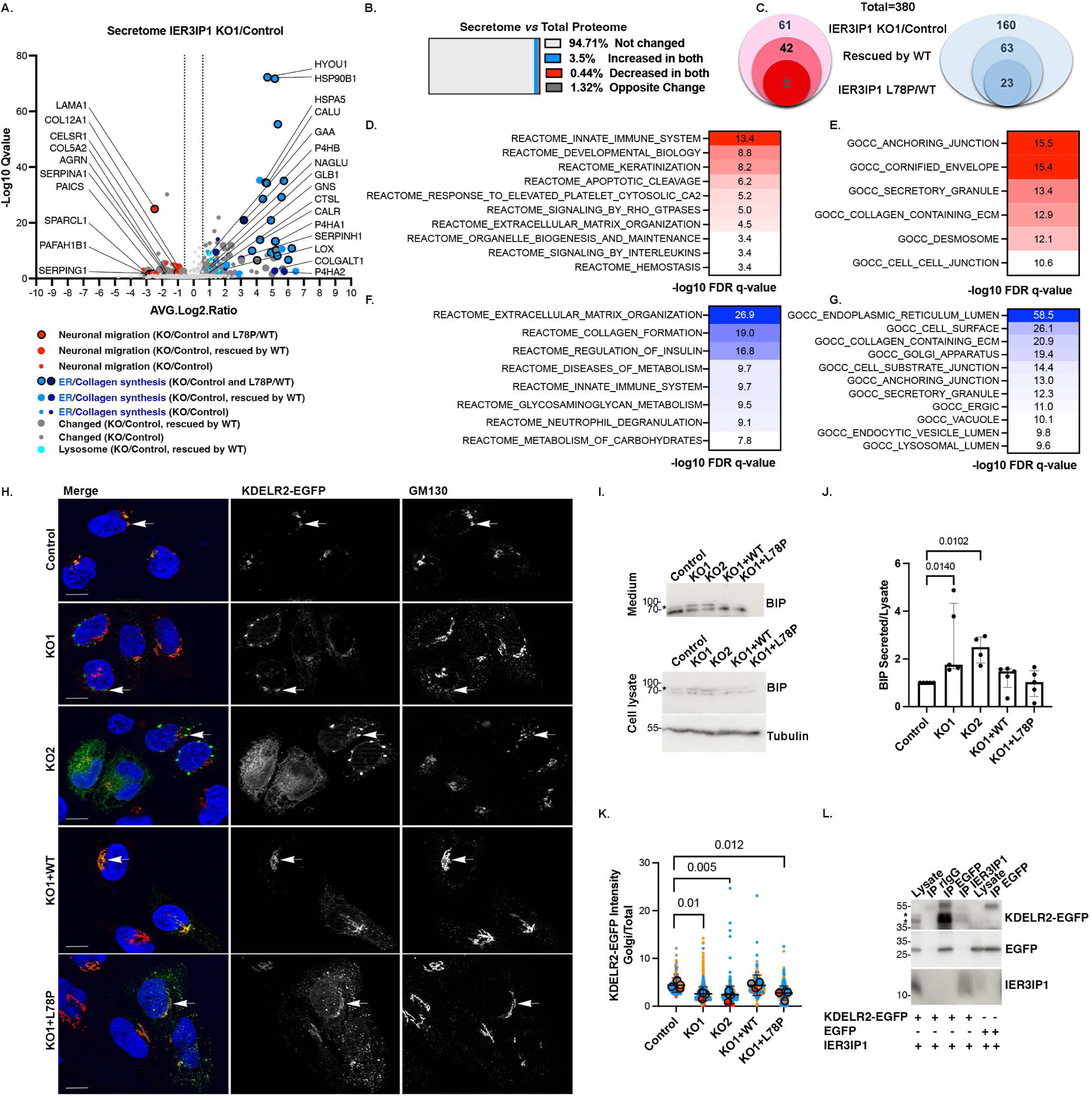
IER3IP1 KO changes the composition of the secretome. **(A)** Volcano plot of proteins differentially secreted by *IER3IP1* KO1 and control cells (Suppl. Table S5). Proteins are color-coded based on their biological roles (legend). Large circles indicate proteins rescued by *IER3IP1* WT and filled large circles proteins whose secretion is significantly different in cells expressing the *IER3IP1* p.L78P mutant *versus IER3IP1* WT (n = 3 for KO1 and WT, and n = 4 for control and p.L78P). **(B)** Percentage of secretome hits unchanged in the total proteome *versus* secretome (grey), increased (blue) or decreased (red) in both conditions, or exhibiting increased secretion but reduced total levels (dark grey) (Suppl. Tables S3, S5). **(C)** Number of proteins whose secretion is decreased (red) or increased (blue) in *IER3IP1* KO1 cells compared to control (outer circle), those rescued by re-expression of the WT (middle circle), and those differentially secreted in cells expressing *IER3IP1* p.L78P compared to WT expressing cells (inner circle) (Suppl Table S5). **(D-G)** GSEA pathway enrichment analysis of proteins whose secretion was decreased (red) or increased (blue) in *IER3IP1* KO1 compared to control cells. **(H, K)** Cells were transfected with KDELR2-EGFP (arrows), fixed, co-labeled with anti-GM130 (red) and Hoechst (blue), and analyzed by fluorescence microscopy. Scale bars, 10 µm. Single Apotome sections are shown. (**K)** Ratios of KDELR2-EGFP fluorescence in the Golgi region and the entire cell were calculated using Fiji. Mean values of independent experiments (large, filled circles), and values for individual cells (small circles) are color-coded for each experiment (n = 4 independent experiments, mean ± SD; One-way Anova with Dunnett’s post-hoc test; n ≥ 195 cells/condition). **(I, J)** BiP immunoprecipitated from cell medium using a specific antibody was analyzed by immunoblotting. **(J)** Ratios between secreted and total cellular BiP levels (n = 4 independent experiments; median and IQR; Kruskal-Wallis with Dunn’s post-hoc test). **(L)** HeLa cells were transfected with the indicated constructs, IER3IP1 and KDELR2-EGFP were immunoprecipitated from lysates using anti-IER3IP1 and anti-EGFP, respectively, and analyzed by immunoblotting with anti-EGFP and anti-IER3IP1 (n = 5 independent experiments).

Interestingly, ∼27% of all the proteins whose secretion was increased in *IER3IP1* KO1 cells were ER resident proteins (Fig. 4A, F, G; Suppl. Table S5). These included ER chaperones such as BiP (termed HSPA5 in the MS data), calreticulin and calnexin, that secure the correct folding and quality control of newly synthesized glycoproteins^25^, and enzymes required for collagen biosynthesis. Re-expression of *IER3IP1* WT rescued the aberrant secretion of most enzymes (Fig. 4A, Suppl. Table S5), and many of them were secreted more by cells expressing the pathogenic mutant p.L78P than by *IER3IP1* WT cells (Suppl. Fig. S5 A, D, E; Suppl. Table S5). Immunoblot analysis of BiP immunoprecipitated from cell culture medium confirmed that cells lacking *IER3IP1* secreted significantly more BiP than control cells, and *IER3IP1* WT re-expression rescued these modifications (Fig. 4I, J). Total cellular levels of BiP were not changed (see also Fig. S3E, F), suggesting abnormal secretion was not caused by an increase of the cellular stress response.

More than half of the ER proteins (55.8%) secreted more in the absence of *IER3IP1* carried ER retention motifs (Suppl. Table S6), suggesting a defect in preventing their KDEL receptor-dependent retrieval to the ER^26^. To test this hypothesis, we analyzed the localization of KDELR1-EGFP, KDELR2-EGFP and KDELR3-EGFP by fluorescence microscopy (Fig. 4H, K; Suppl. Fig. S5 F-I). All three receptors are endogenously expressed in HeLa cells, with KDELR2 displaying the highest mRNA levels^27^. In control cells, all three receptors co-localized to a high degree with the ci-Golgi marker GM130, as expected^27, 28^. In contrast, the fraction of Golgi-localized KDELR2-EGFP was significantly reduced by *IER3IP1* KO, a change rescued by *IER3IP1* WT, but not by *IER3IP1* p.L78P (Fig. 4H, K). The proportion of Golgi localized KDLER1-EGFP was not changed, and that of KDELR3-EGFP was reduced only in *IER3IP1* KO1 cells, compared to controls (Suppl Fig. S5F-I). In addition, IER3IP1 immunoprecipitated KDELR2-EGFP from cell lysates (Fig. 4L). Together, these data suggest IER3IP1 may play a role in the transport of KDELR2 from the ER to the cis-Golgi, ultimately ensuring the retention of various ER-resident proteins.

### Down-regulation of *Ier3ip1* affects neuronal morphology and positioning in the embryonic brain

Our data suggest that loss of IER3IP1 causes changes in secreted and surface-localized proteins involved in neuronal development and migration. However, they do not reveal whether and how such changes may affect corticogenesis. To bridge the gap between our molecular analysis in HeLa cells and neurodevelopmental disorders underlying MEDS1, we next explored the outcomes of *Ier3ip1* depletion *in vivo*. To this end, the lateral cortex of E13.5 mouse embryos was used as a model region. Neural progenitors were targeted by *in-utero* electroporation^29^ with plasmids expressing EGFP and shRNAs against either *Ier3ip1* (Fig. 5B) or luciferase as a control. Their EGFP+ neuronal progeny was analyzed two days later at E15.5 (Fig. 5A). A possible effect of shIer3ip1 on survival was excluded (Fig. 5H, I). Upon leaving the VZ/SVZ, newborn neurons are multipolar and generate several extremely dynamic tangential processes^29, 30^. In the upper part of the IZ, neurons change their polarity and become bipolar while undergoing migration towards the cortical plate^30, 31^. We found that in the lower bins of the IZ, adjacent to the VZ/SVZ, neurons with *Ier3ip1* KD displayed significantly longer neurites compared to controls (Fig. 5A, C), suggesting changes in the neurite extension/retraction cycle. When CP-localized neurons were classified based on their morphologies^32^, the percentage of neurons with complex arborizations was higher in *Ier3ip1* KD cells than in the controls, at the expense of unbranched uni/bipolar cells (Fig. 5D, E). The angles between the leading neurites relative to the orientation of the CP showed a higher variability in *Ier3ip1* KD neurons, where significantly more neurons were tilted at angles of 60°-80° compared to control, where most neurons had orientations between 80°-90° (Fig. 5F, G). In conclusion, the loss of IER3IP1 affects the dynamic transition between subsequent morphological stages, likely by changing the equilibrium of secreted and surface-localized proteins required for the precise coordination of intracellular and extracellular signals^33^.

**Figure 5.**
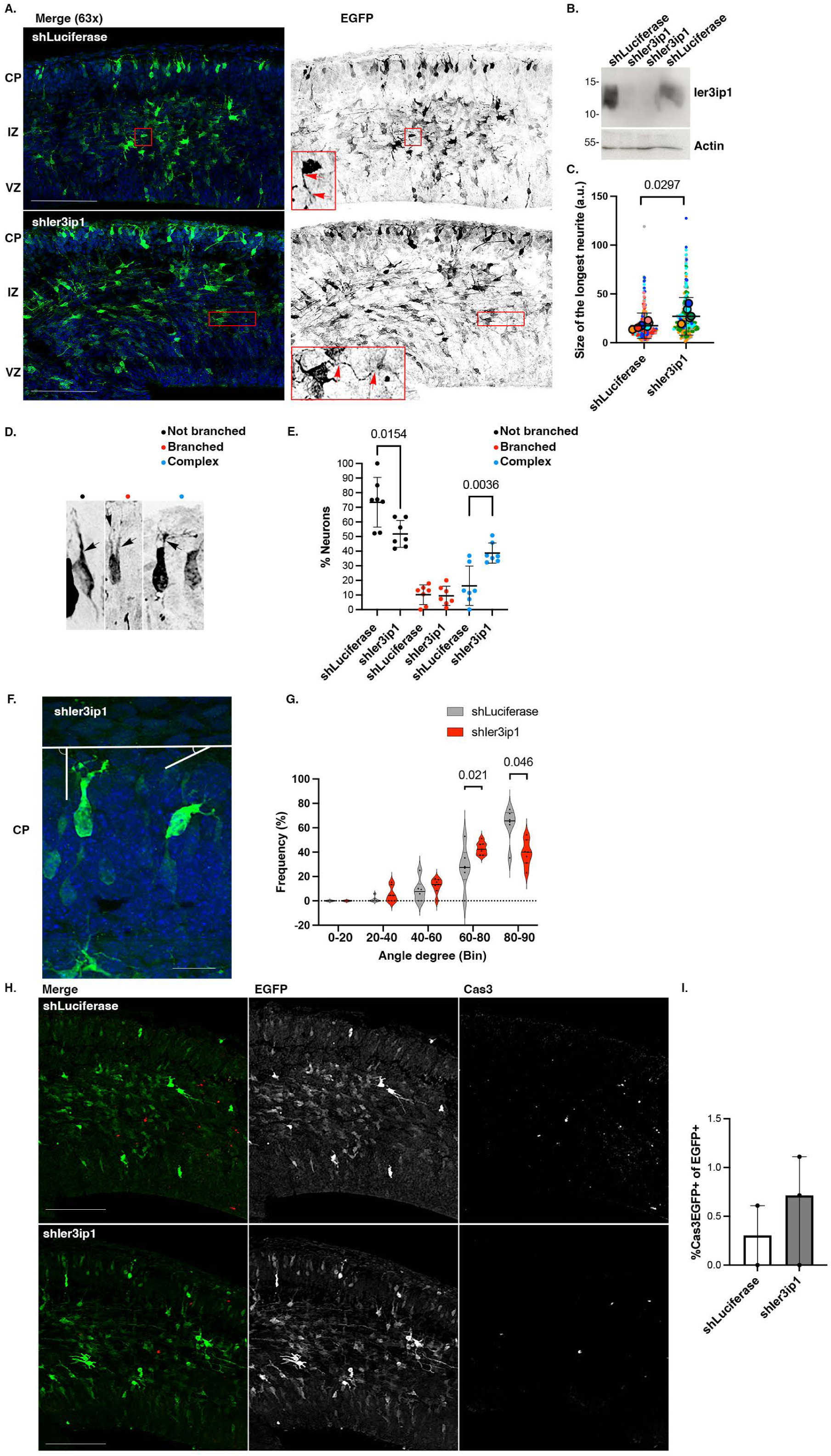
Ier3ip1 knockdown affects neuronal positioning. Brains were electroporated with shRNAs at E13.5 and collected at E15.5. **(A, C)** Coronal sections were fixed and labeled with the indicated antibodies and Hoechst nuclear dye. Right panels show inverted EGFP signal (black). Inserts show selected cells at a higher magnification. VZ, IZ and CP were distinguished based on nuclear density and morphology Hoechst (blue) staining. **(C)** The longest neurite in each analyzed cell (arrows) was measured, and color-coded values for individual cells (small circles) and mean values per brain (circles) are shown (n _shLuciferase_ = 7 brains, 302 cells; n _shIer3ip1_ = 5 brains, 280 cells; mean ± SD; two-tailed Welch’s t-test). **(B)** NIH3T3 cells were transfected with shRNAs, and EGFP+ cells were sorted and analyzed by immunoblotting (n = 3). **(D, E)** EGFP+ neurons in the CP were classified as unbranched uni/bipolar cells (black circles), bipolar branched (red circles) or neurons displaying complex arborizations (blue circles) and quantified (n _shLuciferase_ = 7 brains, 9 sections, 338 EGFP+ cells; n _shIer3ip1_ = 7 brains, 10 sections, 475 cells; mean±SD, two-tailed Welch’s t-test). Scale bars, 100 µm. **(F)** The angles between the leading neurites of EGFP+ neurons and the CP surface were determined, and **(G)** the frequencies (%) of the values within the ranges of values that define the indicated bins are shown as percentages of the total number of values. n _shLuciferase_ = 7 brains, 10 sections, 185 EGFP+ cells; n _shIer3ip1_ = 7 brains, 11 sections, 203 cells; Violin plot with indicated median value (black line) and quartiles (dashed lines), two-tailed Mann-Whitney test. Scale bar, 20 µm. **(H)** Sections were labeled with anti-Cas3 (red) and Hoechst (blue), and **(I)** the percentages of EGFP+ cells additionally positive for Cas3 were calculated. shLuciferase n = 2 brains, 4 sections, 302 EGFP+ cells; shIer3ip1 n = 3 brains, 4 sections, 445 EGFP+ cells, median ± IQR, two-tailed Mann-Whitney test. Scale bars, 100 µm. All images represent maximum projections of Apotome Z-sections.

## Discussion

Here we combined proteomics and functional approaches to characterize changes in membrane trafficking in cells lacking *IER3IP1* or expressing *IER3IP1* mutants that are associated with MEDS1, a fatal syndrome affecting multiple systems and organs, including the brain. Among the *IER3IP1* pathogenic variants, p.T79Δ is not expressed at the protein level, p.L78P negatively affects the organization of the secretory pathway, similarly to *IER3IP1* depletion, whereas p.V21G causes no significant morphological changes, suggesting further investigations are necessary for its characterization. Although previous studies support the importance of IER3IP1 in ER function^10, 14, 15^, a direct role for IER3IP1 in ER to Golgi transport has not been demonstrated in mammalian cells, and its potential cargos have remained unknown. Our data indicate that IER3IP1 is involved in the ER export of a subset of secretory and plasma membrane proteins, some of which play a role in axon pathfinding, neuronal migration, and development. ERGIC53 and KDELR2, two important receptors in the early secretory pathway, are affected by the absence or malfunction of IER3IP1. This suggests a possible molecular mechanism for the changes in surfaceome and secretome, with potential consequences for cellular homeostasis in MEDS1. *In vivo*, the positioning and the morphology of neurons lacking IER3IP1 are modified, suggesting a potential pathomechanism for MEDS1.

### How does IER3IP1 promote the ER exit of specific cargoes?

IER3IP1 controls the export of a subgroup of proteins from the ER, indicating it plays a role in their selective recruitment to ERES. The mechanism could involve IER3IP1 interactions with soluble proteins *via* a luminal domain, as seen for SURF4^34^ and ERGIC53^35^, or the chaperoning of transmembrane domains, as described for the classical cargo receptor cornichon/Erv14p^36^. This function may be performed as a cofactor of cargo receptors like ERGIC53. IER3IP1 could add complexity to the system by increasing the number of cargos or by conferring specificity towards a certain subset of proteins, or perform other roles, such as maintaining trafficking-favorable protein conformations^37^. In agreement with this hypothesis, IER3IP1 binds ERGIC53 and controls both ERGIC53 trafficking and the secretion of SERPINA1 (alpha-1-antitrypsin), a known cargo of ERGIC53^22^. Although ERGIC53 has not been associated with neurological defects but with bleeding disorders, its family member LMAN2L, whose cargoes are not known, is mutated in patients with intellectual disability and epilepsy^38^.

### ER enzymes are not efficiently retrieved to the ER in the absence of IER3IP1

Our data suggest that the role of IER3IP1 in ER-export has implications for another important feature of the early secretory pathway, the retention of ER resident proteins. Rather strikingly, cells lacking *IER3IP1* or expressing *IER3IP1* p.L78P abnormally secrete ER resident proteins with C-terminal KDEL or KDEL-like sequences like BiP, whose localization to the ER requires KDELRs^39^. Three KDELRs (KDELR1-3) traffic between the ER and Golgi via COPII- and COPI-coated carriers^40^, but their specific functions are not well understood. It has been suggested that siRNA-mediated knockdowns of both *KDELR1* and *KDELR2* are necessary to efficiently reduce ER protein retrieval^41^. The interpretation of this study is, however, complicated by the fact that none of the two receptors was fully depleted. In contrast, the knockout of *KDELR1* alone was sufficient for increasing the secretion of an ER resident protein^42^. Only KDELR2 is mislocalized in the absence of IER3IP1, suggesting decreased amounts of KDELR2 on the Golgi are sufficient to cause the abnormal secretion of ER resident enzymes. The importance of KDELR2 is supported by the finding that its mutation causes osteogenesis imperfecta and neurodevelopmental delay^43^. An alternative mechanism that could explain the abnormal chaperone secretion could be similar to the one described for p24^44^. In this case, reduction of p24 diminishes incorporation of its specific cargoes into ERES, allowing the recruitment of cargo normally not secreted.

### Consequences of secretion of ER chaperones

In the case of most ER resident proteins, only small amounts escape the ER and need to be retrieved^45^. Nevertheless, reduced retrieval and increased secretion may ultimately compromise ER quality control^46^, allowing the export of malfunctioning/misfolded proteins. Defective protein folding could result in cargo accumulation, causing the enlargement of the ER cisternae, as observed in the absence of IER3IP1. Similar changes in ER shape have been observed in cells lacking other ER exit regulators such as TANGO1^47^, Sec24^48^, Sec23B^49^, Sec13^50^ or SAR1^51^. Some of the ER chaperones abnormally secreted by IER3IP1-deficient cells are essential for development. A mouse model expressing a secreted BiP mutant has a severe neurological phenotype including microcephaly, defects in neuronal migration, cortical layer organization, and early death^52^. Moreover, SIL1, a nucleotide exchange factor for BiP, is required for maintaining cortical architecture and insulin secretion^53^, and two other enzymes have been directly linked to microcephaly: POFUT1^54^ and MINPP1^55^. Several ER chaperones secreted by IER3IP1-deficient cells (P4HB, P4HA1, P3H1, FKBP10, SERPINH1, CRTAP, COLGALT1, PLOD1, PLOD2, and LOX) are essential for the synthesis of collagens controlling ECM stiffness and cell migration^56^. Furthermore, some ER chaperones have known functions in the extracellular space. Secreted calreticulin mediates fibrinogen-dependent mitogenic activity^57^ or cell spreading^58^, and secreted BiP controls neuronal process growth^59^, processes that could be disturbed by their enhanced secretion, as observed here in the absence of IER3IP1.

Does the secretion of chaperones and the accumulation of selected cargo in the ER cause ER stress? ER stacks appear distended in cells lacking IER3IP1, suggesting a certain level of ER stress, but BiP protein levels are not increased. The ER stress sensors phospho-IRE1 and ATF6 are not activated, as previously observed in pancreatic β-cells^14^. IER3IP1 deficiency-related accumulation of enlarged ER cisternae is likely to activate cellular mechanisms that remove unnecessary membranes for maintaining cellular homeostasis^40^. Lysosomal activity and secretion of lysosomal enzymes were higher in the absence of IER3IP1, disturbances that may also contribute to MEDS1 pathology.

### IER3IP1 controls the composition of the cellular secretome and surfaceome, and thereby neuronal positioning

Several specific cargos exported with the help of IER3IP1 are surface or secreted proteins essential for neuronal function, and their identification may help understand some of the clinical features described for MEDS1. Interestingly, these included an increase of surface and secreted basolateral proteins, which may shift apical-basal polarity during neurogenesis^60^. Neuronal morphological differentiation and migration are precisely coordinated by gradients of diffusible signaling molecules and plasma membrane receptors, surface bound cues and physical properties of the substrate (i.e., ECM stiffness)^56^. Our data suggest that not one, but several factors are changed in cells lacking IER3IP1. For example, UNC5B is a netrin1 repulsive receptor implicated in axon guidance, neurite branching and neuronal migration^61-63^. FGFR3, another receptor essential for development, is enriched in the embryonic neural proliferative zone^64^. Like FGFR2^65^, which is also reduced on the surface of *IER3IP1* KO cells, FGFR3 influences neuronal migration in the embryonic brain^66^. Although cortical development is not affected in Fgfr3-null mice models, microcephaly has been observed in zebrafish lacking fgfr3^67, 68^. Moreover, a potentially inactivating mutation in the catalytic domain of human FGFR3 causes developmental delay and, in some patients, microcephaly^69^. Two other examples of receptors reduced on the surface of *IER3IP1* KO cells are TGFBR3 and ACVR2B, components of the TGF beta signaling pathway, which is essential for neuronal differentiation and nervous system development^70^. In contrast, secreted ligands such as BMP2, BMP6, SEMA4B, or the surface-localized neuropilin 1, a semaphorin receptor essential for neuronal guidance^71^ are increased in the absence of IER3IP1.

IER3IP1 is also critical for the transport of ECM components, including collagen COL5A2 or the collagen-binding SPARCL1, whose reduction may affect collagen fibrillogenesis. Another example is LAMA1, whose extracellular levels are reduced in *IER3IP1* KO cells, in agreement with observations in *IER3IP1* KO brain organoids^15^. LAMA1 is a major component of basal membranes, whose mutations affect neuronal development^72^. Interestingly, LAMA1 and transient axonal glycoprotein 1 (TAG-1), an adhesion molecule, are essential for the polarization of multipolar cells in the IZ^73^. In contrast, a subset of integrins, which are laminin receptors, is upregulated on the surface of *IER3IP1* KO1 cells. Together, these data suggest that laminin/integrin signaling, essential for progenitor proliferation and migration^74^, as well as for neuronal migration^75^, is disrupted in the absence of IER3IP1.

How does deficiency of IER3IP1 cause microcephaly? *In vivo*, Ier3ip1 depletion alters neuronal morphology in the developing cortex. Neurite morphology is essential for establishing neuronal connectivity and changes have been linked to neurodevelopmental disorders^76^. For instance, microcephaly has been associated with mutations in *RAB3GAP/RAB18*^77, 78^, *ARFGEF2*^79^, *ARF1*^80, 81^, kinesin *KIF2A*^2^, or dynein heavy chain^82^, all affecting both membrane trafficking^83^ and neuronal morphogenesis. Future studies will investigate if, in addition to neuronal polarity, *IER3IP1* mutation also affect neural progenitor identity and/or migration.

In summary, we found that IER3IP1 deletion affects the surface levels and/or the secretion of multiple molecules implicated in neuronal development, suggesting that not a single factor, but the different composition of the neuronal membrane and its environment are responsible for the observed phenotype.

### Limitations of the Study

Our research underscores the mislocalization of key proteins in neurodevelopment, migration, and axon pathfinding due to IER3IP1 deficiency, influencing neuronal morphology. However, our study has certain limitations. We lack a detailed understanding of the specific impact of individual mislocalized proteins on the observed phenotype. Future studies should unravel the distinct roles of these proteins in the context of IER3IP1 deficiency. Additional research is needed to explore IER3IP1’s role in different cell types in the cortex and during various developmental stages. While our study touches upon neuronal morphology, further exploration is needed to understand the effects of IER3IP1 on progenitor proliferation, neurogenesis and migration. The distinction between cell-extrinsic and cell-intrinsic roles of mislocalized secreted factors remains unclear and should be addressed in future investigations. Our study does not delve into certain aspects of MEDS1, such as juvenile diabetes implications and the specific role of malfunctioning IER3IP1.

In conclusion, while our study provides valuable insights into IER3IP1’s role in neurodevelopment and microcephaly, addressing these limitations through further research will enhance our understanding of the intricate molecular mechanisms involved.

## Star Methods

| **Antibodies** | **Source** | **Application and dilution**<br>(IB, immunoblot; IF, immunofluorescence; IP, immunoprecipitation) |
| --- | --- | --- |
| P62/SQSTM1 clone 2C11 | H00008878-MO1, Abnova | IB:1000 |
| ATF6 | ab227830, Abcam | IB:1000 |
| V5 Tag monoclonal antibody SV5-Pk1 | R960-25, Thermo Fischer Scientific | IB 1:1000, IF 1:100 |
| GFP mouse IgG<sub>1</sub>K (clones 7.1 and 13.1) | 1181446001, Roche, Sigma-Aldrich, Merck | IB 1:1000, IP 2 μl |
| FGF Receptor 3 (C51F2) | 4574, Cell Signaling Technology | IB 1:1000 |
| ST6GAL1 | AF5924, R&D Systems | IF 1:100 |
| LC3B | NB100-2220, Novus Biologicals, Bio-Techne | IB 1:1000 |
| IER3IP1 | HPA010027, Atlas Antibodies, Sigma-Aldrich | IB 1:500, IF 1:40 |
| alpha-tubulin clone DM1A | T6199, Sigma-Aldrich | IB 1:1000 |
| beta-actin | ab8227, Abcam | IB 1:1000 |
| Flag clone M2 | F1804, Sigma-Aldrich | IB 1:1000, IF 1:200 |
| BIP | 3177, Cell Signaling Technology | IB 1:1000 |
| recombinant anti-IRE1 (phospho S724) EPR5253 | ab 124945, Abcam | IB 1:1000 |
| IRE-1a (14C10) | 3294, Cell Signaling Technology | IB 1:1000 |
| Cas3 clone C92-605 | 559565, BD Biosciences | IF 1:600 |
| GFP-1020 IgY | Aves Labs, 2BScientific | 1:400 |
| LAMP1 human CD107a, clone H4A3 | 555798, BD Biosciences | IF 1:100 |
| calnexin | ADI-SPA-860D, Enzo Life Science | IF 1:50 |
| Sec31A clone 32 | 612350, BD Biosciences | IB, 1:1000, IF 1:200 |
| Sec16A | ab70722, Abcam | IB 1:1000, IF 1:200 |
| ERGIC53 cl B9 | sc-271517, Santa Cruz Biotechnology | IF 1:200 |
| GM130 clone 35 | 610822, BD Biosciences | IF 1:200 |
| Normal Rabbit IgG | 2729, Cell Signaling Technology |  |
| HRP-anti-Rabbit IgG(H+L) | W4011, Promega GmbH | IB: 1:5000 |
| HRP-anti-Mouse IgG(H+L) | W4021, Promega GmbH | IB: 1:2000 |
| HRP-Donkey anti-goat IgG | sc-2020, Santa Cruz Biotechnology | IB: 1:2000 |
| Donkey anti-Mouse IgG (H+L) 555 | A-31570, Invitrogen™, Thermo Fisher Scientific | IF: 1:500 |
| Goat anti-Mouse IgG (H+L) 488 | A-11001, Invitrogen™, Thermo Fisher Scientific | IF: 1:500 |
| Goat anti-Mouse IgG (H+L) 555 | A-21422, Invitrogen™, Thermo Fisher Scientific | IF: 1:500 |
| Goat anti-Mouse IgG (H+L) 594 | A-11005, Invitrogen™, Thermo Fisher Scientific | IF: 1:500 |
| Rabbit IgG (H+L) | A-11034, Invitrogen™, Thermo Fisher Scientific | IF: 1:500 |
| Goat anti-Rabbit IgG (H+L) 555 | A-21428, Invitrogen™, Thermo Fisher Scientific | IF: 1:500 |
| Goat anti-Rabbit IgG (H+L) 647 | A-21244, Invitrogen™, Thermo Fisher Scientific | IF: 1:500 |
| Goat anti-Rat IgG (H+L) 488 | A-11006, Invitrogen™, Thermo Fisher Scientific | IF: 1:500 |
| Goat anti-Rat IgG (H+L) 555 | A-21434, Invitrogen™, Thermo Fisher Scientific | IF: 1:500 |
| Donkey anti-Sheep IgG (H+L) 555 | A-21436, Invitrogen™, Thermo Fisher Scientific | IF: 1:500 |
| Donkey anti-Goat IgG (H+L) | A-11055, Invitrogen™, Thermo Fisher Scientific | IF: 1:500 |
| Donkey anti-Chicken IgY (H+L) 488 | A78948, Invitrogen™, Thermo Fisher Scientific | IF: 1:500 |

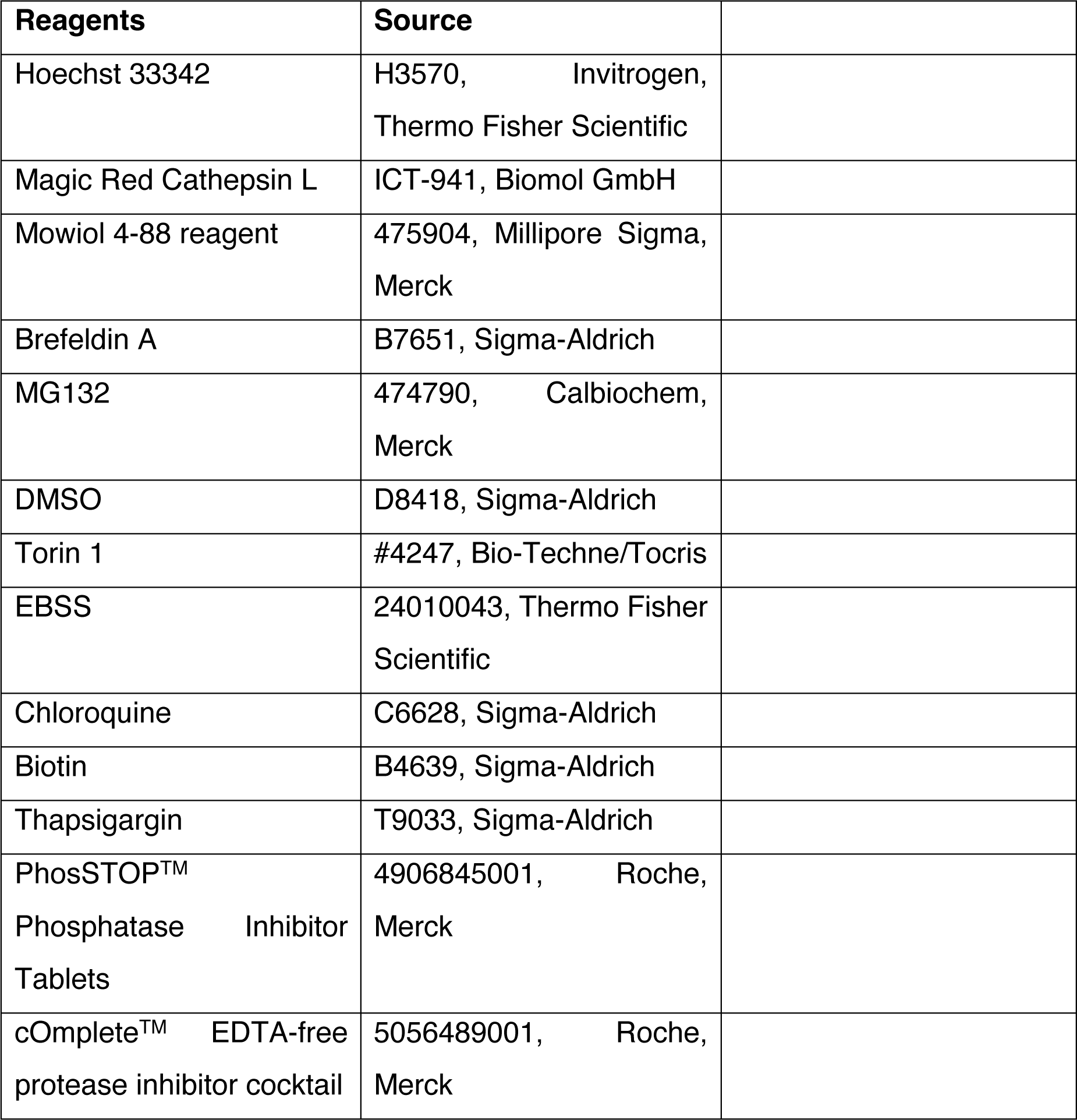
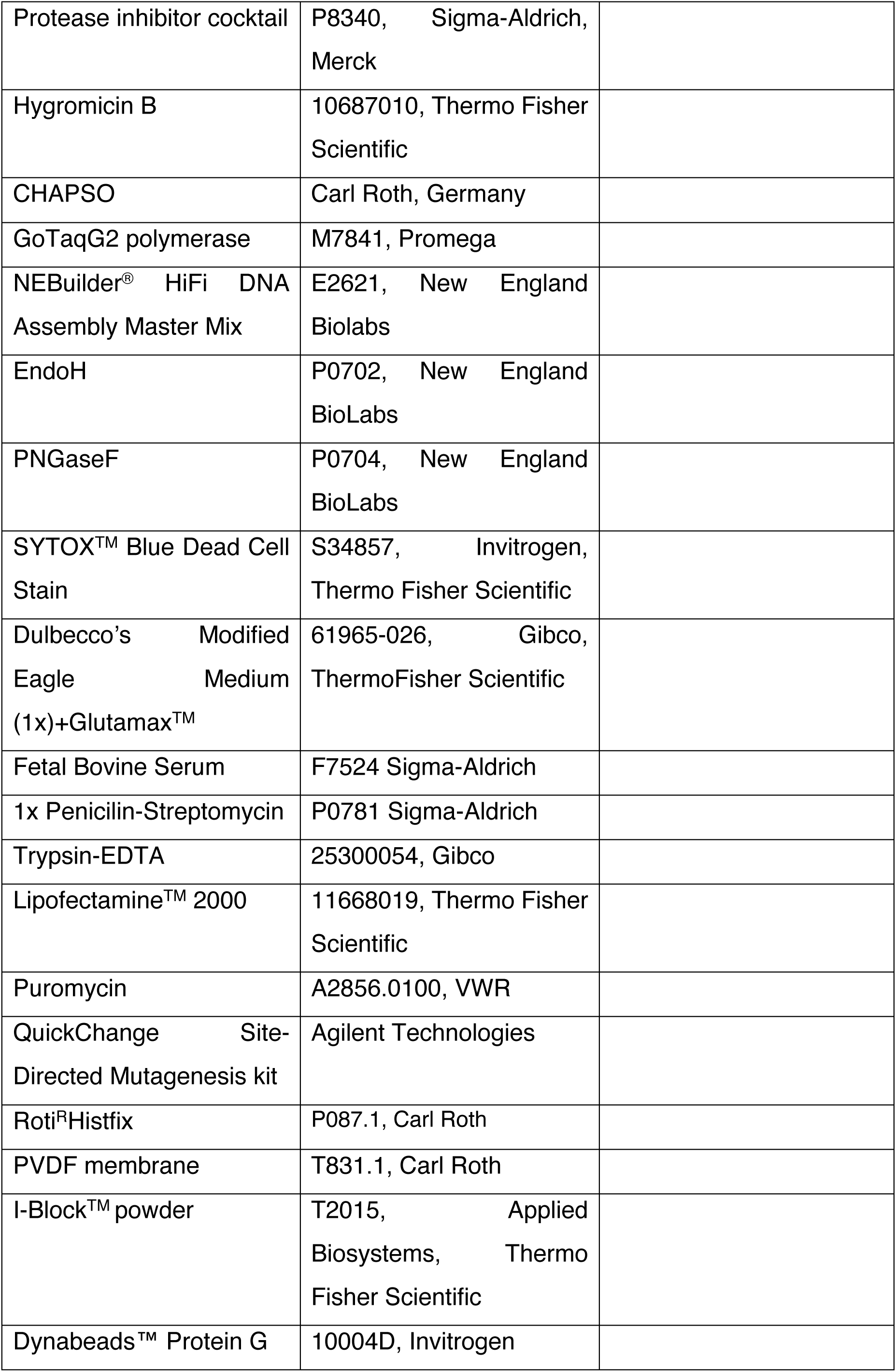
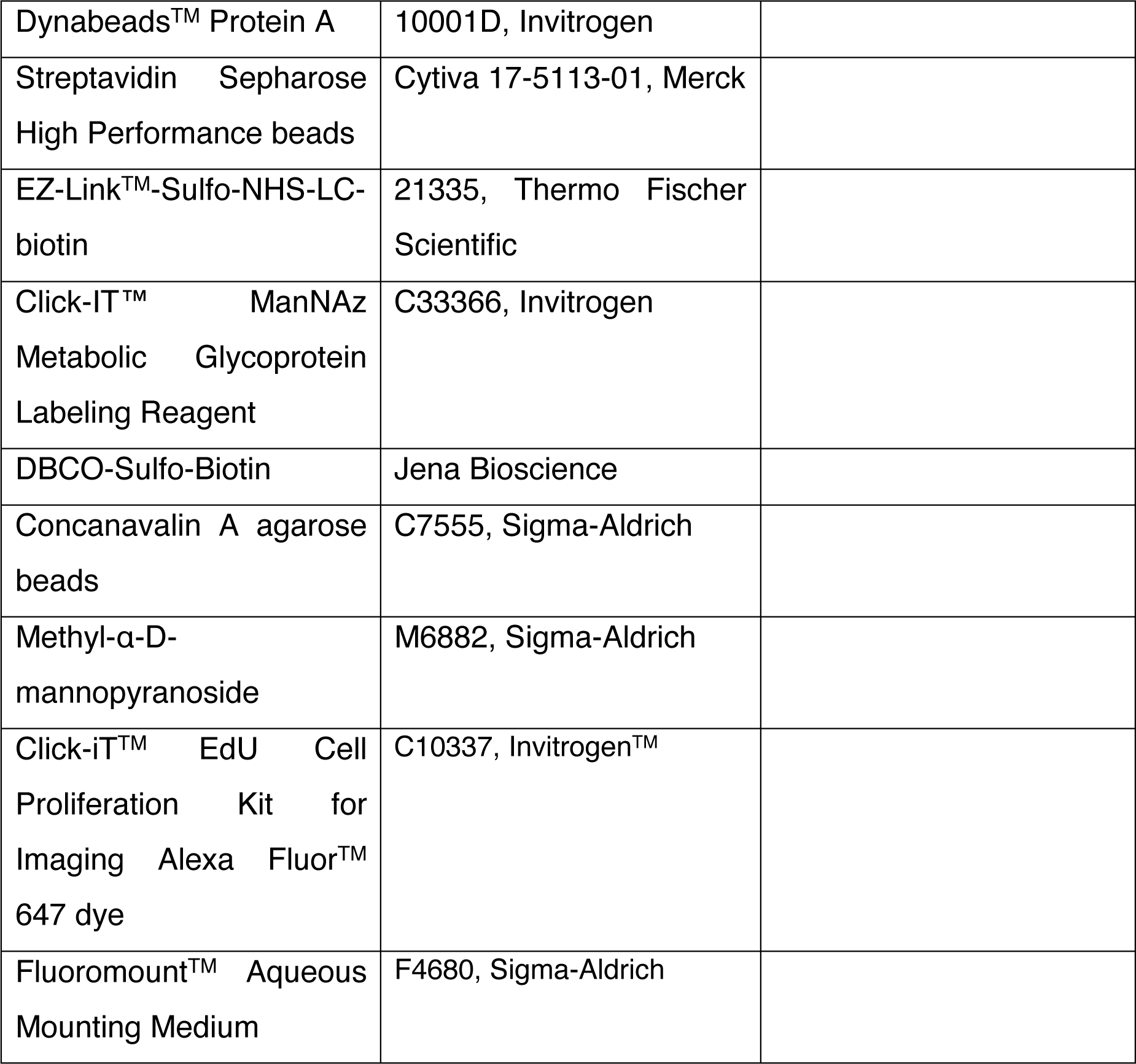

| Reagents | Source |
| --- | --- |
| Hoechst 33342 | H3570, Invitrogen, Thermo Fisher Scientific |
| Magic Red Cathepsin L | ICT-941, Biomol GmbH |
| Mowiol 4-88 reagent | 475904, Millipore Sigma, Merck |
| Brefeldin A | B7651, Sigma-Aldrich |
| MG132 | 474790, Calbiochem, Merck |
| DMSO | D8418, Sigma-Aldrich |
| Torin 1 | #4247, Bio-Techne/Tocris |
| EBSS | 24010043, Thermo Fisher Scientific |
| Chloroquine | C6628, Sigma-Aldrich |
| Biotin | B4639, Sigma-Aldrich |
| Thapsigargin | T9033, Sigma-Aldrich |
| PhosSTOP™<br>Phosphatase Inhibitor<br>Tablets | 4906845001, Roche, Merck |
| cOmplete™ EDTA-free<br>protease inhibitor cocktail | 5056489001, Roche, Merck |
| Protease inhibitor cocktail | P8340, Sigma-Aldrich, Merck |
| Hygromycin B | 10687010, Thermo Fisher Scientific |
| CHAPSO | Carl Roth, Germany |
| GoTaqG2 polymerase | M7841, Promega |
| NEBuilder<sup>®</sup> HiFi DNA Assembly Master Mix | E2621, New England Biolabs |
| EndoH | P0702, New England BioLabs |
| PNGaseF | P0704, New England BioLabs |
| SYTOX<sup>™</sup> Blue Dead Cell Stain | S34857, Invitrogen, Thermo Fisher Scientific |
| Dulbecco's Modified Eagle Medium (1x)+Glutamax<sup>™</sup> | 61965-026, Gibco, ThermoFisher Scientific |
| Fetal Bovine Serum | F7524 Sigma-Aldrich |
| 1x Penicilin-Streptomycin | P0781 Sigma-Aldrich |
| Trypsin-EDTA | 25300054, Gibco |
| Lipofectamine<sup>™</sup> 2000 | 11668019, Thermo Fisher Scientific |
| Puromycin | A2856.0100, VWR |
| QuickChange Site-Directed Mutagenesis kit | Agilent Technologies |
| Roti<sup>®</sup>Histfix | P087.1, Carl Roth |
| PVDF membrane | T831.1, Carl Roth |
| I-Block<sup>™</sup> powder | T2015, Applied Biosystems, Thermo Fisher Scientific |
| Dynabeads<sup>™</sup> Protein G | 10004D, Invitrogen |

|  |  |  |
| --- | --- | --- |
| Dynabeads™ Protein A | 10001D, Invitrogen |  |
| Streptavidin Sepharose High Performance beads | Cytiva 17-5113-01, Merck |  |
| EZ-Link™-Sulfo-NHS-LC-biotin | 21335, Thermo Fischer Scientific |  |
| Click-IT™ ManNAz Metabolic Glycoprotein Labeling Reagent | C33366, Invitrogen |  |
| DBCO-Sulfo-Biotin | Jena Bioscience |  |
| Concanavalin A agarose beads | C7555, Sigma-Aldrich |  |
| Methyl-α-D-mannopyranoside | M6882, Sigma-Aldrich | |
| Click-iT™ EdU Cell Proliferation Kit for Imaging Alexa Fluor™ 647 dye | C10337, Invitrogen™ |  |
| Fluoromount™ Aqueous Mounting Medium | F4680, Sigma-Aldrich |  |

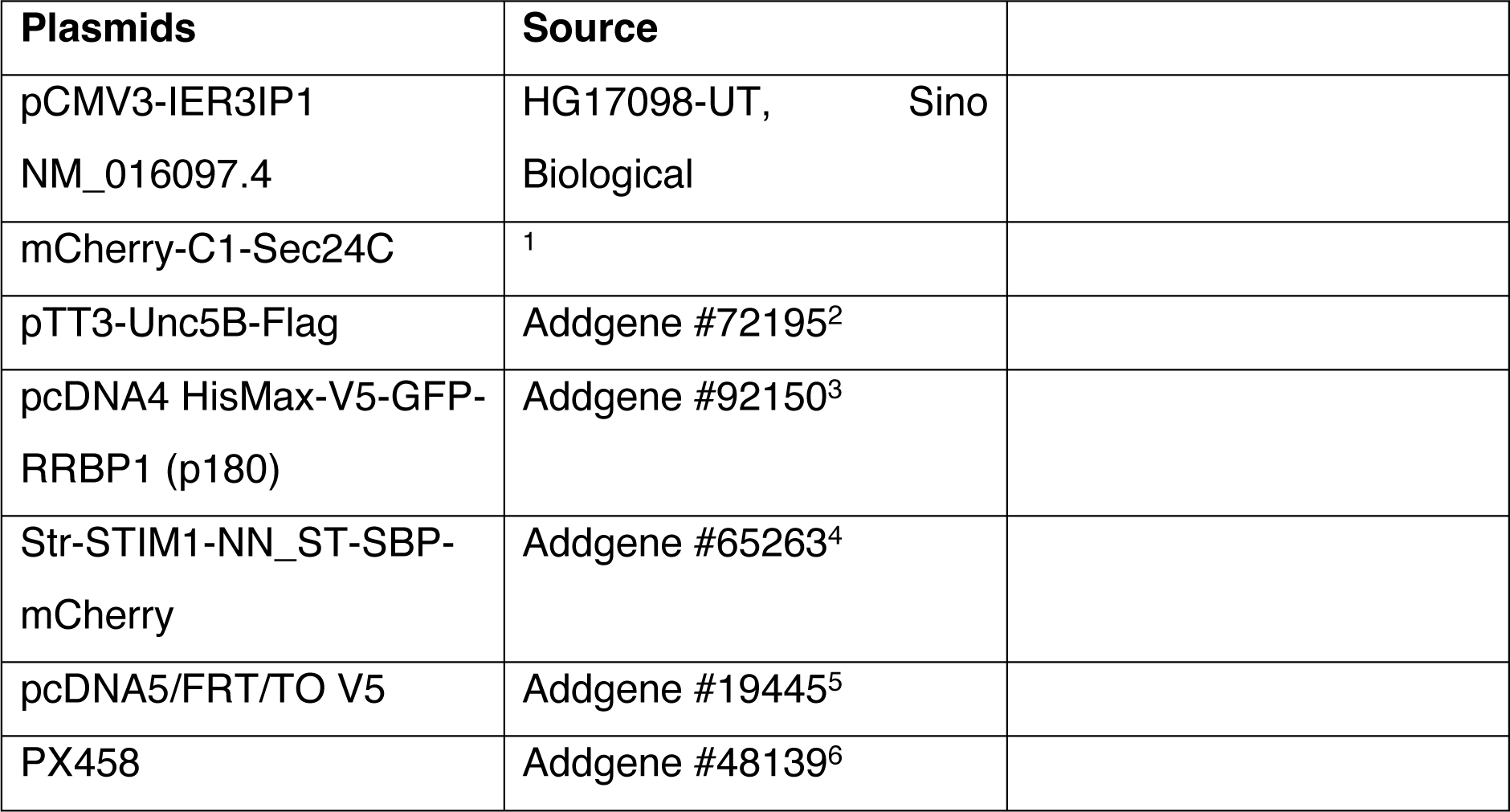
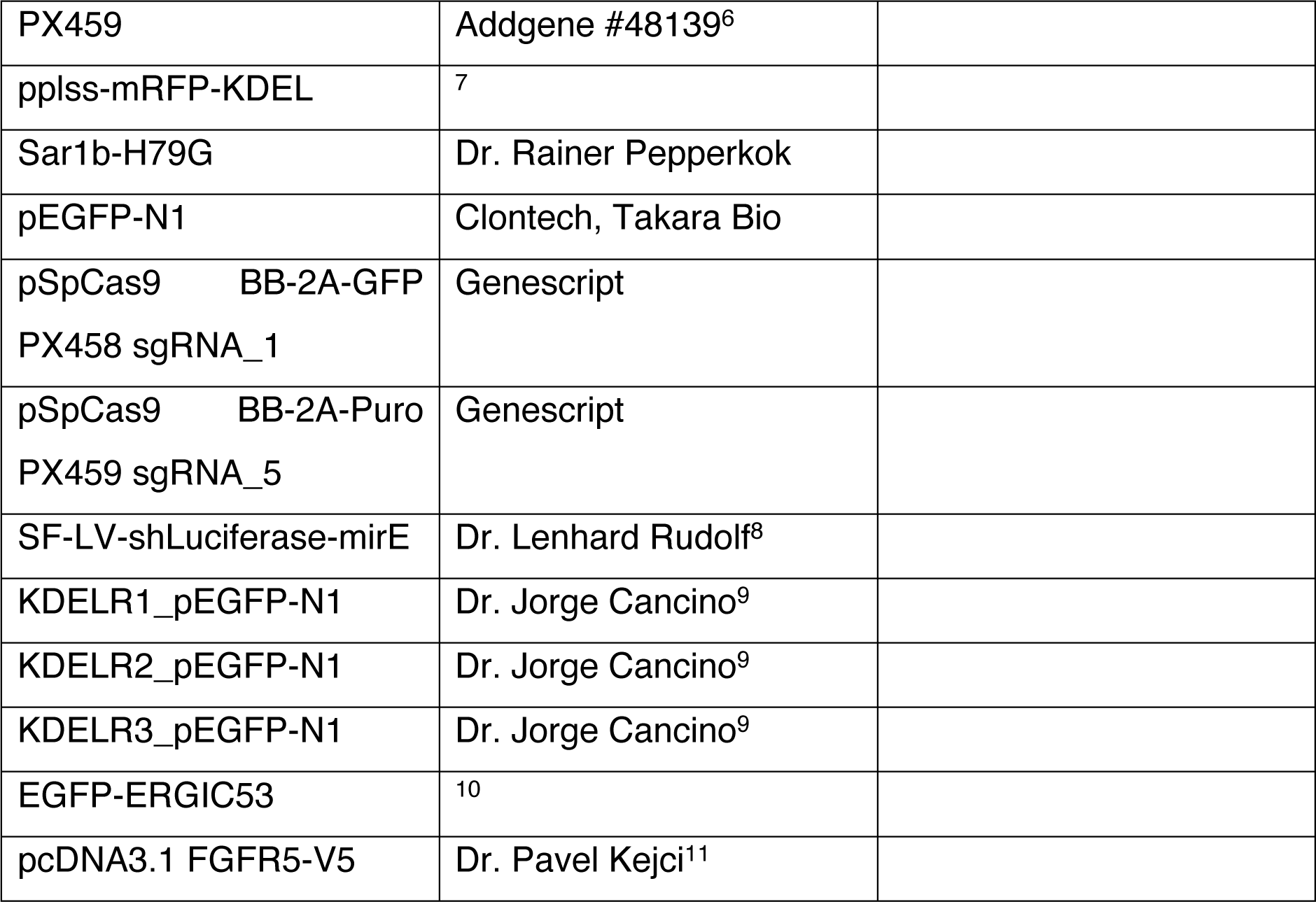

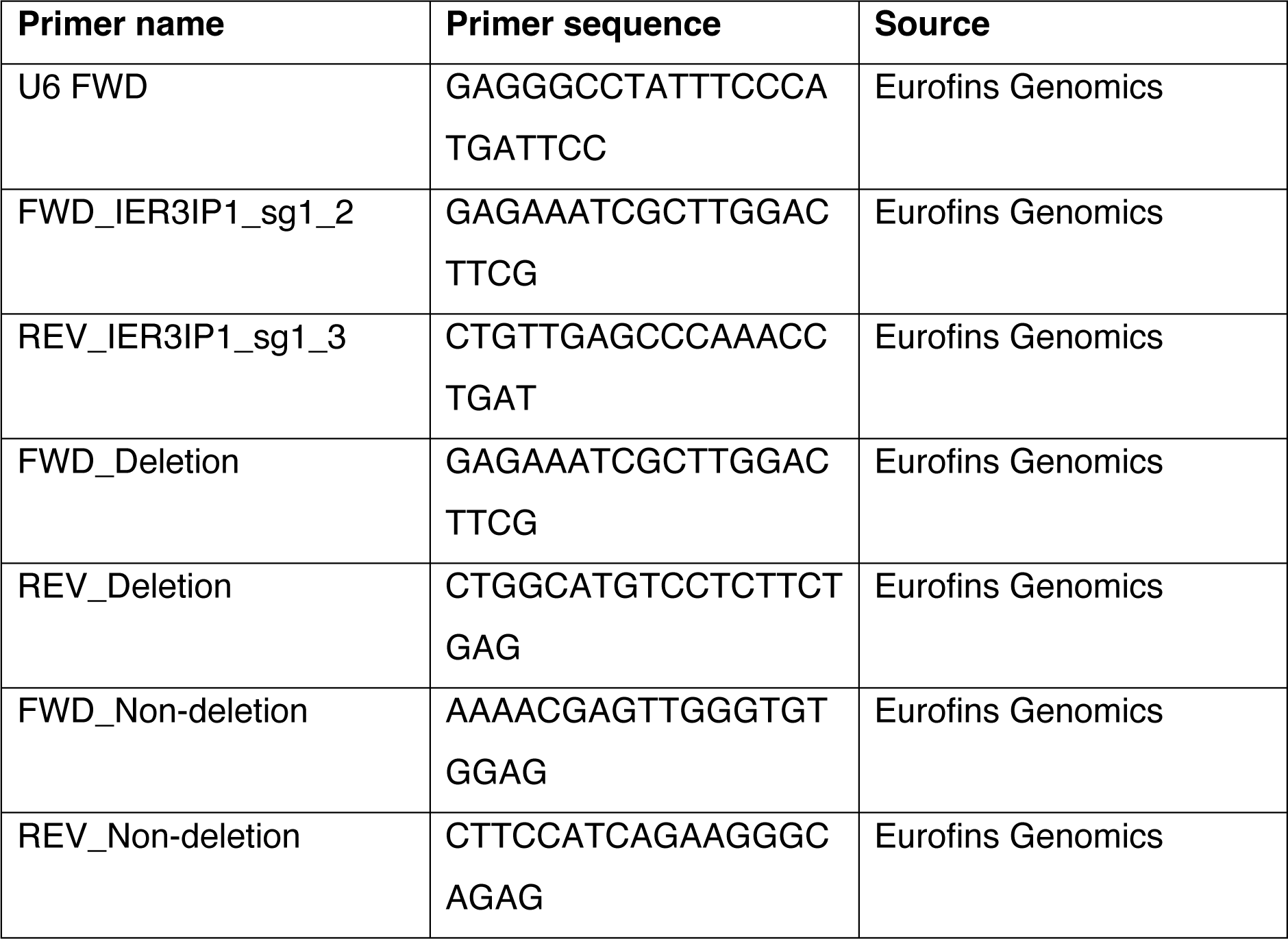
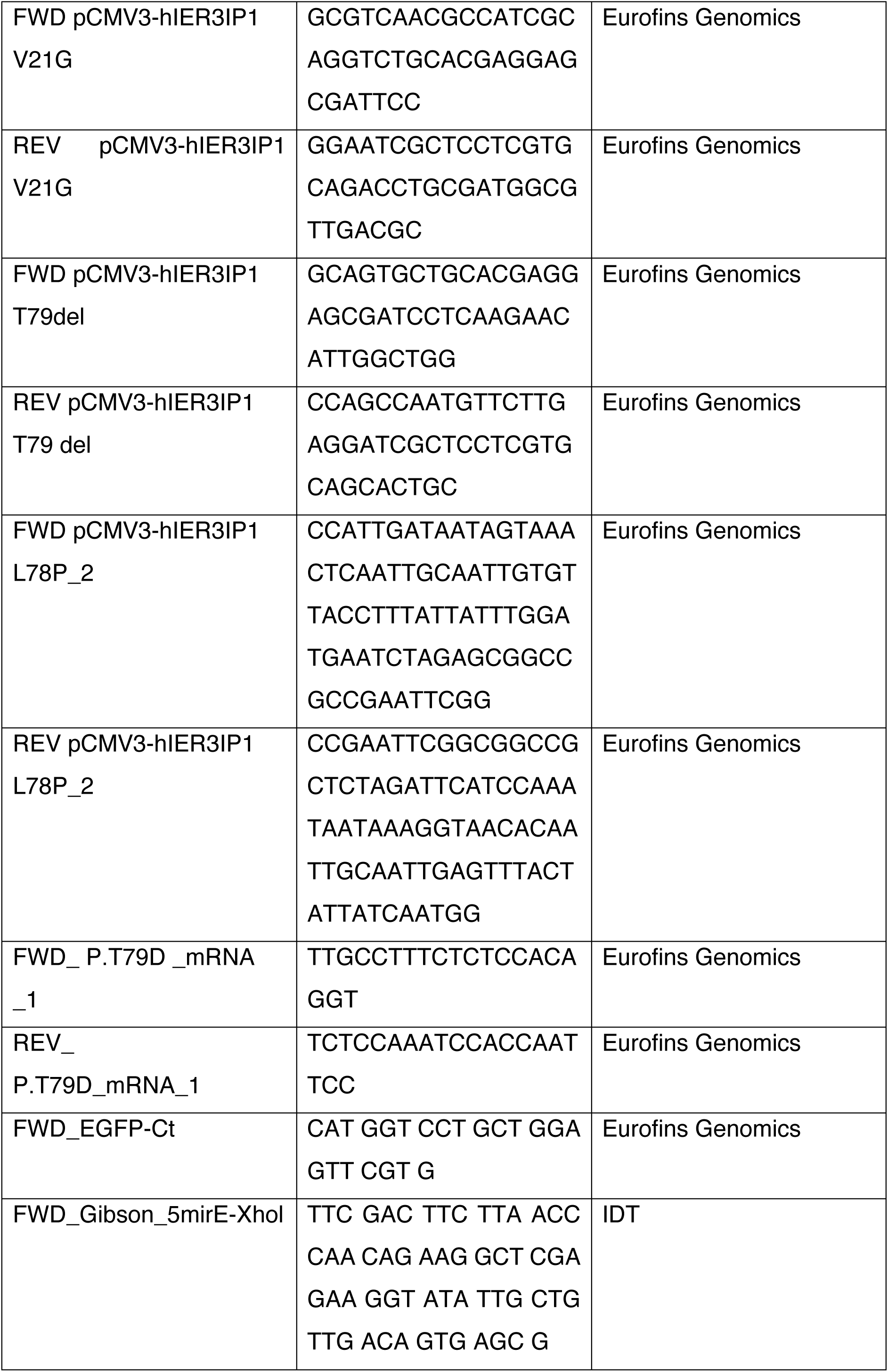
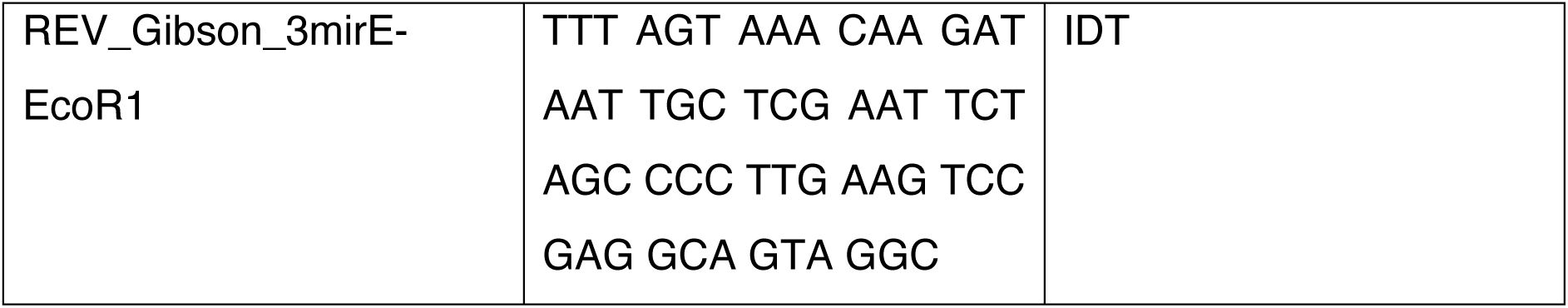

### Cell culture

HeLa Kyoto cells were grown in Dulbecco’s Modified Eagle Medium (1x)+Glutamax^TM^ (61965-026, Gibco, ThermoFisher Scientific) with 10% Fetal Bovine Serum (F7524 Sigma-Aldrich) and 1x Penicilin-Streptomycin (P0781 Sigma-Aldrich), and incubated at 37°C and 5% CO_2_. Cells were detached using Trypsin-EDTA (25300054, Gibco) and washed with 1xPBS.

### Transfection

For transient transfection, 24 h after seeding, cells were incubated with DNA and Lipofectamine^TM^ 2000 (11668019, Thermo Fisher Scientific), according to manufacturer’s protocols. To obtain cell lines stably expressing IER3IP1 variants, KO1 IER3IP1 cells were plated in 6-well plates and transfected with 2.4 µg DNA and 7 µl of Lipofectamine per well. After 24 h, 200 µg/ml hygromicin B were added to each well, and 72 h later, 10 cells/ml were transferred to 10 cm plates. After 10-14 days, monoclonal cell colonies were picked and transferred to 24-well plates. Clones were analyzed for IER3IP1 expression and localization by immunoblot and immunofluorescence microscopy, and suitable clones were further grown with 200 µg/ml hygromicin B.

### CRISPR-Cas9 knockout

sgRNAs were designed using CHOPCHOP ^84^: sgRNA_1 CGCCATCGCAGTGCTGCACG (targeting exon 1) and sgRNA_5 ATTGAGTTTACTATTATCAA (targeting exon 3). The two sgRNAs were cloned in pSpCas9 BB-2A-GFP PX458 and pSpCas9 BB-2A-Puro PX459 sgRNA_5, respectively, by Genescript, and analyzed by sequencing with the U6 FWD primer. HeLa cells were transfected with Lipofectamine 2000, and pSpCas9 BB-2A-GFP PX458 sgRNA_1 and pSpCas9 BB-2A-Puro PX459 sgRNA_5 to knockout IER3IP1, or the control vectors PX458 and PX459 to obtain a control cell line. After 1 day, we sorted the top 5% GFP-positive cells using fluorescence activated cell sorting (FACS). Sorted cells were resuspended in medium supplemented with 1.5 µg/ml Puromycin (A2856.0100, VWR), and plated in 6-well plates (100,000 cells/well). After 24 h, cells were washed twice with 1xPBS, grown in complete medium for 1 more day, and then transferred to either 10 cm plates (100 cells/plate) or to 96 well plates (0.5 cells in 200 µl per well), and grown in complete medium for 2 weeks, until colonies formed. Clonal colonies were selected and transferred to 24-well plates and then to 6-well plates.

The presence of INDEL mutations was analyzed by sequencing (see Fig. 1). After genomic DNA isolation, the targeted region was amplified by PCR using specific primers and GoTaq polymerase (Promega), according to manufacturer’s protocol. The entire IER3IP1 gene was amplified with FWD_Deletion and REV_Deletion primers, and a region downstream of the sgRNA_1 targeted region with FWD_Non-deletion and REV_Non-deletion primers. The region targeted by sg_RNA_1 was amplified with the FWD_IER3IP1_sg1_2 and REV_IER3IP1_sg1_3 primers, and Sanger sequences of these amplicons were analyzed using the Inference of CRISPR Edits tool (ICE; ice.synthesgo.com). The KO2 clone was selected from a total of 10 clones as having the highest (93%) ICE KO score. The KO1 clone had a large deletion that comprised the entire IER3IP1 gene. To confirm the KO of IER3IP1 we analyzed clones by immunoblot and immunofluorescence. Both the KO1 and the KO2 clones displayed morphologies and growth rates comparable to control (Cas9-transfected) cells. Images were created using the SnapGene software (Fig. 1A-D).

### Site-Directed Mutagenesis

was done using the QuickChange Site-Directed Mutagenesis kit (Agilent Technologies, CA, USA), according to the suggested protocol. The primers used were, for pCMV3-IER3IP1 V21G: FWD pCMV3-hIER3IP1 V21G, REV pCMV3-hIER3IP1 V21G, for pCMV3-IER3IP1 L78P: FWD pCMV3-hIER3IP1 L78P_2, REV pCMV3-hIER3IP1 L78P_2, and for pCMV3-IER3IP1 T79: FWD pCMV3-hIER3IP1 T79del, REV pCMV3-hIER3IP1 T79 del.

### Immunocytochemistry and microscopy

Cells grown on No.1 glass coverslips (0111550, Paul Marienfeld GmbH, Germany) were treated as indicated, then washed with 1xPBS, fixed with Roti^R^Histfix (P087.1, Carl Roth, Germany) (20 min at room temperature), permeabilized with 0.2% Triton X-100 in PBS (5 min), and blocked with a solution containing 1 ml FCS, 1 g BSA, 0.1g fish gelatin in 100 ml 1xPBS for 20 min. Cells were then incubated with the primary antibodies diluted in blocking solution (1 h, room temperature), washed 3 x 5 min with PBS, then labeled with secondary antibodies (20 min, room temperature), washed 3 x 5 min with PBS, and mounted on microscope slides (AA00000112E01MNZ10, Epredia, Germany) using Mowiol 4-88 with Hoechst 33342 (1:1000). Images were acquired using an Imager Z.2 equipped with an Apotome.2 and Axiocam 702, with a Plan-Apochromat 63x/1.4 Oil DIC M27 objective (Carl Zeiss AG, Germany).

For live cell imaging, cells were plated in 8-well Chambered Coverglass w/non-removable wells No. 1 (155411, Thermo Scientific^TM^ Nunc ^TM^ Lab-Tek ^TM^, Thermo Fisher Scientific). Images were acquired using an Axio Observer Z1/7 equipped with an Apotome.2 and Axiocam 702 Mono, with a Plan-Apochromat 63x/1.40 Oil DIC M27 objective (Carl Zeiss AG).

For high-throughput imaging, cells grown in Eppendorf Cell Imaging Plate, 96-well (Eppendorf AG, Germany) were either incubated with Magic Red^TM^ Cathepsin L (1:250) and 5 µg/ml Hoechst 33342 (60 min, 37°C), or fixed and labeled with anti-LAMP1 and Hoechst 33342. Cells were imaged with an ImageXpress Micro Confocal microscope equipped with a 40x, NA=0.95 CFI Plan Apo Lambda air immersion objective, temperature, and humidity control (Molecular Devices, CA), Air/CO2 gas mixer from OKOLAB, Italy) and MetaXpress High-Content Image Acquisition and Analysis v.6.7.2.290 (64-bit) software. For LAMP1, Z-stacks were acquired and MaxProjections were analyzed.

### RUSH assay

Cells grown on coverslips were transfected with Str-STIM1-NN_ST-SBP-mCherry (ST-mCherry)^4^ and Lipofectamine. After 24 h, 40 µM biotin was added into the medium, and cells were incubated for the indicated times at 37°C, then fixed and immunostained. The mean fluorescence intensity of the mCherry signal was measured in the Golgi, GM130-positive, region and in the entire cell, using Fiji^13^. Ratios between the Golgi and the total cellular average intensities were calculated using Excel.

### ER shape analysis

Cells grown in 8-well chambers were transfected with GFP-p180 and Lipofectamine 2000, and live-cell imaged after 1 day. Fiji was used to measure, for individual cells, the GFP-p180-positive area/cell (after thresholding using the Fiji “Moments” algorithm) and the total cell area, and the ratio between the two areas was calculated per cell.

### To measure the number of objects

per cell for the different markers analyzed, as well as the co-localization between GM130 and ST6GAL1 (object-based Pearson’s correlation coefficient), images were processed with Fiji, and object thresholding, detection and measurement per image were performed with Cell Profiler (www.cellprofiler.org)^14^.

### IncuCyte growth curve analysis

Cells were plated in Nunc™ MicroWell™ 96-Well, Nunclon Delta-Treated, Flat-Bottom Microplate (167008 Thermo Fisher Scientific) (5000 cells/well) and imaged every 3 h for the indicated times, using the IncuCyte S3 Live-Cell Analysis System (Sartorius, Germany) with an IncuCyte S3 microscope (4647 Essen BioScience) and a 10x objective. In each well, images were acquired at four distinct positions. Cell confluence (i.e., the area occupied by cells) was measured, and the average confluence per well was calculated with the IncuCyte S3 software.

### Transmission electron microscopy

Cells were fixed with Karnovsky fixative (2% paraformaldehyde, 3% glutaraldehyde in 0.1M Cacodylate buffer, pH 7.3), for 3 h at room temperature, then overnight at 4°C. After 5 x 15 min washes with Cacodylate buffer, cells were incubated (protected from light) with 2% osmium tetroxide + 1% potassium hexacyanidoferrate (II) in 0.1 M Cacodylate buffer, at 4°C for 2 h, and then washed 3 x 15 min with Cacodylate buffer. Fixed cells were scrapped and centrifuged (2000 x rpm using a table centrifuge, room temperature), then embedded in 3% agar, and cut into small pieces with a razor blade. Using a tissue processor (Leica, Germany) these cuts were washed 3 x 15 min with Cacodylate buffer, 3 x 15min with distilled water, dehydrated by incubating with acetone with increasing concentrations: 30, 50, 70, 90, 95 and 3x 100% (30 min each), and stained with 1% uranyl acetate in 50% acetone. Sample infiltration was performed with epoxy resin (Glycid ether 100, #21045 SERVA Electrophoresis GmbH, Germany): a mixture of acetone:resin 3:1 (45 min), 1:1 (45 min), and 1:3 (45 min), then 3 x 2 h pure epoxy resin, and 1 x 2h epoxy resin + accelerator (Benzyl Dimethylamine or BDMA). Samples were embedded in flat moulds, allowed to polymerize at 60°C for 48 h, then trimmed with a Reichert UltraTrim (Leica). Semithin sections (0.5 µm) were labeled with Azure staining^85^, then ultrathin sections (55 nm) made with an ultramicrotome “Reichert Ultracut S” (Leica) were placed onto copper slot grids coated with a Formvar/Carbon layer. Images were obtained with a transmission electron microscope JEM 1400 (JEOL, MA) with a CCD camera ‘Orius SC 1000A’ (GATAN, CA), an acceleration voltage of 80 kV, and GATAN MICROSCOPY SUITE 2.31.734.0 software.

### RNA Expression analysis

Cells were scraped and collected in PBS, then centrifuged (1500xg, 2 min, 4°C). Pellets were lysed, RNA was extracted using the NucleoSpin RNA, mini kit for RNA purification (740955.50, Machery-Nagel, Germany). cDNA was synthesized using the qScript cDNA Synthesis Kit (95047, Quanta BioSciences, MA), as indicated by the manufacturer, and analyzed by PCR using the FWD_ P.T79D _mRNA _1 and REV_ P.T79D_mRNA_1 primers.

### SDS-PAGE and Immunoblotting

Cells grown to confluence in 6-well or 10 cm plates were washed with ice-cold PBS, detached with a cell scraper in lysis buffer (50 mM Tris-HCl pH 7.6, 150 mM NaCl, 2 mM EDTA, 1% NP-40 and protease inhibitors) or CHAPSO buffer (1% CHAPSO, 150 mM NaCl, 5 mM EDTA, 50 mM Tris pH 7.6 and protease inhibitors), lysed on ice for 30 min, then centrifuged (500xg, 5 min, 4°C). Supernatants were collected and proteins were denatured by boiling at 90°C for 5 min with sample buffer. Proteins were separated by SDS-PAGE, transferred to Immobilon-P PVDF membranes (T831.1, Carl Roth). Page Ruler^TM^ Plus Pre-stained Protein Ladder (26619, Thermo Fisher Scientific) was used as protein size standard. Membranes were incubated in blocking solution (1g I-Block^TM^ powder (T2015, Applied Biosystems, Thermo Fisher Scientific), 0.1% Tween-20 (9127.2, Carl Roth) in 500 ml of 1xPBS)), with primary antibodies diluted in blocking solution (1 h at room temperature or overnight at 4°C), washed 3 x 5 min with 1xTBS-Tween, incubated with secondary antibodies (30 min, room temperature) and washed 3 x 5 min with 1xTBS-Tween.

#### Immunoprecipitation

Equal numbers of cells were plated in 10 cm plates and transfected the next day with the indicated plasmids and Lipofectamine 2000. After 24 h, cells were detached, collected in CHAPSO buffer, lyzed 30 min on ice, then centrifuged 13 000xg, 40 min at 4°C. Supernatants were transferred to low binding Protein LoBind Tube 1.5. ml (022431081, Eppendorf), and incubated with the indicated antibodies (10 µl for anti-IER3IP1, 5 µl for the other antibodies), in the cold room with rotation, overnight. Next day, pre-washed 80 µl of Dynabeads™ Protein G (10004D, Invitrogen) were added to each tube, incubated in the cold room with rotation, for 30 min. Beads were then washed 5 x 1 ml lysis buffer, transferred to a new tube before the last wash, then resuspended in sample buffer, and analyzed by SDS-PAGE and immunoblot.

#### Immunoprecipitation from culture medium

Equal number of cells were plated in 4x 10 cm plates for each condition. After 24 h, medium was replaced with 5 ml of fresh medium, and cells were incubated at 37°C for 2 more days. Media from plates corresponding to the same experimental condition were combined, and volumes were adjusted to correspond to equal numbers of cells. After addition of protease inhibitors, media were filtered through a 0.45 µm Rotilabo PVDF filter (P667.1, Carl Roth), loaded onto a Vivaspin 20, 30 kDa Diafilter (VS2022, Sartorius AG) and concentrated by centrifugation (4000xg, 4°C, 80 min) to a volume of 0.5 ml of retentate. Retentates were transferred to low binding tubes, 2 µl of anti-BiP were added to each tube, followed by an incubation at 4°C, with rotation, overnight. The next day, 100 µl of prewashed Dynabeads^TM^ Protein A (10001D, Invitrogen) were added to each tube, followed by a 30 min incubation at 4°C, with rotation. Beads were further washed 5 x 1 ml lysis buffer and resuspended in Sample buffer. Cells were lysed in lysis buffer (50 mM Tris-HCl pH 7.6, 150 mM NaCl, 2 mM EDTA, 1% NP-40 and protease inhibitors) and analyzed by immunoblot.

#### Glycosylation assays

HeLa cells transfected with Unc5B-FLAG and Lipofectamine™ 2000 were treated with EndoH (P0702, New England BioLabs) or PNGaseF (P0704, New England BioLabs), at 37C°C for 16 h, as indicated by the manufacturer, then analyzed by immunoblot.

#### Autophagy analysis^15^

Cells grown in 6-well plates were incubated with growth medium containing DMSO or 50 µg/ml Chloroquine (Sigma-Aldrich, C6628), or 500 nM Torin 1 (Bio-Techne/Tocris) in EBSS starvation medium, for 5 h, then lysed and analyzed by SDS-PAGE and immunoblot.

#### ER stress analysis

Cells grown in 6-well plates were treated with 1 µM thapsigargin or an equal volume of control DMSO for the indicated times, lysed and analyzed by immunoblot.

#### Acetylation of beads

Streptavidin Sepharose High Performance beads (Cytiva 17-5113-01, Merck) were equilibrated in PBS and lysine- acetylated using 20 mM sulpho-NHS-acetate for 1 hour at room temperature. Sulpho-NHS-acetate was quenched by adding 1 M Tris pH 7.5 and beads were extensively washed with PBS.

#### Surface biotinylation

Cells were grown in 10 cm plates for 2 days, then processed on ice. Each plate was washed with 3 x 5 min x 5 ml of ice-cold PBS with Ca^2+^ and Mg^2+^, then incubated with 3 ml of 0.5 mg/ml of EZ-Link^TM^-Sulfo-NHS-LC-biotin (21335, Thermo Fischer Scientific) for 30 min, washed 4 x 15 min x 5 ml of 20 mM glycin in PBS, then with 5 ml PBS with Ca^2+^ and Mg^2+^, and collected in lysis buffer (50 mM Tris-HCl pH 7.6, 150 mM NaCl, 2 mM EDTA, 1% NP-40 and protease inhibitors). Lysates were incubated on ice for 30 min, centrifuged at 6000 x g, 4°C, for 6 min. For each sample, 160 µl of acetylated beads (50% suspension) were equilibrated in lysis buffer before adding them to lysates overnight, at 4°C, rotating at 15 rpm. For immunoblot analysis, beads were washed 1 x 1 ml Buffer 1 (50 mM Tris HCl pH 7.6, 325 mM NaCl, 2 mM EDTA, 0.2% NP-40), 5 x 1 ml Buffer 2 (50 mM Tris-HCl pH 7.6, 150 mM NaCl, 2 mM EDTA, 1% NP-40, 1% SDS) and 5 x 1ml Buffer 3 (50 mM Tris-HCl pH 7.6, 150 mM NaCl, 2 mM EDTA), then resuspended in sample buffer with 3 mM biotin, and analyzed by immunoblot. For mass spectrometry analysis, beads were washed with 1 ml of Buffer 1, 4 ml of Buffer 2, and 5 ml of Buffer 3. Beads were finally washed 5 times with 600 µl Wash Buffer 2 (50 mM Ammonium Bicarbonate/AmBic pH 8.0).

#### Secretome protein enrichment with click sugars^21^

To label glycoproteins, cells (5 x 10 cm plates/condition) were incubated with medium with 62.5 µM Click-IT™ ManNAz Metabolic Glycoprotein Labeling Reagent (C33366, Invitrogen) for 48 h. Cell growth medium was supplemented with EDTA-free protease inhibitors and filtered (0.45 µm). Volumes were adjusted to correspond to equal cell numbers and loaded onto Vivaspin 20, 30 kDa Diafilter and concentrated by centrifugation (4000 x g, 4°C, 80 min) to 0.5 ml, then washed 2 x 15 ml PBS at 4°C. The volume of each sample was adjusted to 1 ml and incubated with 100 µM DBCO-Sulfo-Biotin (Jena Bioscience, Germany) overnight at 4 °C, then washed 3 x 15 ml PBS. Concanavalin A agarose beads (C7555, Sigma-Aldrich) were equilibrated by washing 2 x 1 ml Binding buffer (5 mM MgCl_2_, 5 mM MnCl_2_, 5 mM CaCl_2_, 500 mM NaCl in 20 mM Tris-HCL pH 7.5). 300 µl of beads were added to each sample together with 1 ml Binding buffer in low binding tubes, then incubated at 4°C, with rotation, for 2 h. Beads were washed 3 x 1 ml Binding buffer, and proteins were eluted twice with 500 µl Elution buffer (500 mM Methyl-α-D-mannopyranoside (M6882, Sigma-Aldrich), 10 mM EDTA, 20 mM Tris-HCL pH 7.5) for 30 min at 4 °C. Combined eluates were passed through Pierce^TM^ Spin Columns (69705, Thermo Fisher Scientific) and then divided into two low binding tubes. 0.5 ml of 2% SDS in PBS and 300 µl of acetylated streptavidin beads were added to each tube, and samples were incubated at 4 °C with rotation at 15 rpm, overnight. Afterwards, samples were centrifuged (2000 x g, 4 °C, 5 min), and beads were resuspended in PBS, transferred to a Spin Column, washed 1 x 1 ml PBS and 3 x 1 ml Wash Buffer 1 (30 mM AmBic, 3 M Urea), and finally 5 x 600 µl Wash Buffer 2 (50 mM AmBic pH 8.0).

### Mass spectrometry analysis

#### On-bead digest for pulldowns

Beads were transferred to a new tube using Wash Buffer 2, centrifuged at 2000 x g for 5 min and the supernatant discarded. Beads were resuspended in 200 μl Wash Buffer 2 and 1 µg LysC added. After an incubation overnight at 37°C, peptides were eluted two times with 150 µl Wash Buffer 2. Elutions were further digested with 0.5 µg trypsin for 3 h at 37°C. The day after, digests were acidified by the addition of trifluoroacidic acid (TFA) to a final concentration of 1% (v/v), then desalted with Waters Oasis® HLB µElution Plate 30 µm (Waters Corporation, MA) under a soft vacuum, following the manufacturer instructions. Briefly, columns were conditioned with 3 x 100 µL solvent B (80% (v/v) acetonitrile; 0.05% (v/v) formic acid) and equilibrated with 3 x 100 µL solvent A (0.05% (v/v) formic acid in Milli-Q water). Samples were loaded, washed 3 times with 100 µL solvent A, and then eluted into 0.2 mL PCR tubes with solvent B. Samples were dried with a speed vacuum centrifuge and stored at −20 °C until LC-MS analysis.

#### Data Acquisition and Processing for DIA Samples

Prior to analysis, samples were reconstituted in MS Buffer (5% acetonitrile, 95% Milli-Q water, with 0.1% formic acid) and spiked with iRT peptides (Biognosys AG, Switzerland). Peptides were separated in trap/elute mode using the nanoAcquity MClass Ultra-High Performance Liquid Chromatography system (Waters, Waters Corporation, Milford, MA) equipped with a trapping (nanoAcquity Symmetry C18, 5 μm, 180 μm × 20 mm) and an analytical column (nanoAcquity BEH C18, 1.7 μm, 75 μm × 250 mm). Solvent A was water and 0.1% formic acid, and solvent B was acetonitrile and 0.1% formic acid. 1 µl of the sample (∼1 μg on column) were loaded with a constant flow of solvent A at 5 μl/min onto the trapping column. Trapping time was 6 min. Peptides were eluted via the analytical column with a constant flow of 0.3 μl/min. During the elution, the percentage of solvent B increased in a nonlinear fashion from 0–40% in 120 min. Total run time was 145 min, including equilibration and conditioning. The LC was coupled to an Orbitrap Exploris 480 (Thermo Fisher Scientific, Bremen, Germany) using the Proxeon nanospray source. The peptides were introduced into the mass spectrometer via a Pico-Tip Emitter 360-μm outer diameter × 20-μm inner diameter, 10-μm tip (New Objective, MA), heated at 300°C, and a spray voltage of 2.2 kV was applied. The capillary temperature was set at 300°C. The radio frequency ion funnel was set to 30%. For DIA data acquisition, full scan mass spectrometry (MS) spectra with mass range 350–1650 m/z were acquired in profile mode in the Orbitrap with resolution of 120,000 FWHM. The default charge state was set to 3+. The filling time was set at maximum of 60 ms with limitation of 3 × 10^6^ ions. DIA scans were acquired with 40 mass window segments of differing widths across the MS1 mass range. Higher collisional dissociation fragmentation (stepped normalized collision energy; 25, 27.5, and 30%) was applied and MS/MS spectra were acquired with a resolution of 30,000 FWHM with a fixed first mass of 200 m/z after accumulation of 3 × 10^6^ ions or after filling time of 35 ms (whichever occurred first). Data were acquired in profile mode. For data acquisition and processing of the raw data Xcalibur 4.3 (Thermo Fisher Scientific) and Tune version 2.0 were used.

#### Data Analysis for DIA Samples

Spectronaut (v. 16, Biognosys AG). DIA data were then uploaded and searched against this spectral library in Spectronaut. Data were searched with the following modifications: Oxidation (M), Acetyl (Protein N-term). A maximum of 2 missed cleavages for trypsin and 5 variable modifications were allowed. The identifications were filtered to satisfy FDR of 1% on peptide and protein level. Relative quantification was performed in Spectronaut for each paired comparison using the replicate samples from each condition. The data (candidate table) and data reports (protein quantities) were then exported. To select significant proteins, a log2FC cutoff of 0.58 and a Q_value_ <0.05 were defined.

### In utero electroporation

#### shRNA cloning

97 bp shRNA ultramers were designed for the reference sequence NM_025409.3 using SPLASH RNA^16^. The top three SPLASH shRNAs were synthesized by IDT:

**NM_025409.3_235_v2**:TGCTGTTGACAGTGAGCGACAGGAATTAAATCTCAACTAATAGTGAAGCCACAGATGTATTAGTTGAGATTTAATTCCTGGTGCCTACTGCCT CGGA;

**NM_025409.3_1373_v2**:TGCTGTTGACAGTGAGCGCAAAGTTGACTTTTCATATTAATAGTGAAGCCACAGATGTATTAATATGAAAAGTCAACTTTATGCCTACTGCCTC GGA;

**NM_025409.3_1340_v2**:TGCTGTTGACAGTGAGCGCTGGCATCTTTCTGTATAGAAATAGTGAAGCCACAGATGTATTTCTATACAGAAAGATGCCATTGCCTACTGCC TCGGA.

For each shRNA, DNA fragments generated by PCR using GoTaqG2 polymerase (Promega) and the primers FWD_Gibson_5mirE-Xhol and REV_Gibson_3mirE-EcoR1 were cloned in the SF-LV-shRNA-mirE vector under the spleen-focus forming promoter (SFFV)^8, 17^, using NEBuilder^®^ HiFi DNA Assembly Master Mix (New England Biolabs, MA), and 3 clones/shRNA were sequenced using the FWD_EGFP-Ct primer (Eurofins Genomics). To test knockdown efficiency, NIH3T3, grown in the same conditions as HeLa cells, cells were transfected with one selected clone per shRNA and lipofectamine. EGFP-positive cells were sorted after 3 days and analyzed by immunoblot and immunofluorescence microscopy. shRNA NM_025409.3_235_v2 showed the highest knockdown efficiency and was selected for further experiments. A construct expressing shLuciferase (AGGAATTATAATGCTTATCTA) was used as a negative control.

In utero electroporation was performed as described in Mestres and Calegari, 2022 (https://www.biorxiv.org/content/10.1101/2022.12.22.521648v1.full). Shortly, pregnant (E13.5) C57BL/6J wildtype (Janvier) mice were anesthetized with isoflurane, and ∼ 2 µl of 2 mg/ml of plasmid and 0.01% Fast Green were microinjected in the embryonic brains (lateral ventricle), followed by electroporation (6 pulses of 30 V and 5 ms) with an electroporator ECM830 Square Wave Electroporation System (BTX, MA). Surgical wounds were sutured after uterus repositioning, and mice were transferred to a new box when awake. After one day, 1 dose of 1 mg/kg of EdU was administered intraperitoneally, and, after one more day, animals were sacrificed by cervical dislocation. Animal procedures followed TVV16/2018 and were approved by local authorities.

Dissected brains were fixed in 4% paraformaldehyde overnight, then 40 µm coronal sections were obtained using a Microm HM 650 V Vibratome (Thermo Fisher Scientific). Sections were mounted on Superfrost^TM^Plus Adhesion Microscope Slides (J1800AMNZ, Epredia, Germany), dried 30 min at room temperature and washed with PBS (15 min), followed by microwave antigen retrieval with 10 mM sodium citrate buffer (pH 6.0) and washed 3 x 10 min with PBS. Sections were incubated with blocking buffer (5% normal goat serum, 1% BSA, 4% Triton X-100 in PBS) for 2 h at room temperature in a humid chamber, then with primary antibodies diluted in blocking solution (overnight at 4°C). Next day, sections were washed 3 x 15 min with PBS, incubated with secondary antibodies diluted (1:1000) in blocking solution (2 h, room temperature), washed 3 x 15 min with PBS, incubated with Click-iT^TM^ EdU Cell Proliferation Kit for Imaging Alexa Fluor^TM^ 647 dye (Invitrogen^TM^) for 30 min, washed for 5 min with PBS, then incubated with Hoechst (1:10 000) for 30 min and washed 3 x 15 min with PBS. Sections were then stained with 0.1% Sudan Black B in 70% ethanol for 20 min, washed 3 x 5min with PBS-Tween (0,02%), then embedded in Fluoromount^TM^ Aqueous Mounting Medium (#F4680, Sigma-Aldrich) and covered with #1 coverslips (H878 and 1870.2, Carl Roth). Images were acquired using an Imager Z.2 equipped with an Apotome.2 and Axiocam 702, with a Plan-Apochromat 63x/1.4 Oil DIC M27 or a 20x/0.8 M27 objective (Carl Zeiss AG). Maximal intensity projections of serial Z-stacks were analyzed using Fiji.

To characterize neuronal morphologies in the CP, EGFP-positive cells were classified as (1) unbranched uni/bipolar, when the leading process was not branched, (2) branched bipolar, with a branched leading process or (3) complex, with a leading process displaying a more complex arborization^32^. For multipolar cells located in the IZ, for each cell, the longest detectable neurite assignable to a cell body was measured using the “Measure” function in Fiji.

**The model of IER3IP1 transmembrane domains** and the localization of the pathogenic mutations was designed using AlphaFold^12^ and created with BioRender.com (Fig. 1J).

**Statistical analysis** was done with GraphPad Prism version 10.0.0 for Windows or 9.0.2 for Mac (GraphPad Software, Boston, MA, www.graphpad.com). The specific statistical test and number of biological replicates are always indicated in the respective figure legend. Only significant p-values are shown.

## Acknowledgements

This work was supported by a grant from the Deutsche Forschungsgemeinschaft (KA1751/8-1). IM and FC were supported by the CRTD and TU-Dresden. We are grateful to Dr. Jorge Cancino (San Sebastian University, Chile) for the KDELR constructs, Dr. Pavel Kejci (Masaryk University, Brno, Czechia) for the FGFR3-V5 and FGFR2-V5 plasmids, Dr. Rainer Pepperkok for Sar1b-H79G, and to Dr. Lenhard Rudolf, FLI, Jena, Germany for SFLV-shLuciferase-mirE, to Dr. Stefan Lichtenhaler, Dr. Karsten Nalbach and Dr. Christian Behrends for advice regarding secretome analysis, Dr. Torsten Kroll for help with high-throughput imaging, and Katrin Buder for support with TEM. We would also like to thank the Imaging and FACS facilities, FLI Jena, Jana Hamann and Daniela Reichenbach for technical support.

## Author contributions

C.K. contributed to project design, data analysis, wrote the manuscript.

M.A. contributed to project design, performed experiments, data analysis, wrote the manuscript.

F. B. contributed to shRNA cloning, brain sectioning and processing of embryonic brain sections, and secretome isolation.

C.V. contributed to the sectioning and the processing of embryonic brain sections.

T.D. and E.C. contributed to MS sample preparation and performed the MS analysis.

M. L. performed *in utero* electroporation and brain dissection, contributed to *in vivo* data analysis.

F.C. provided the means to perform *in utero* electroporation and contributed to *in vivo* data analysis.

## Supplementary Figures

**Supplementary Figure S1. Characterization of IER3IP1 localization and of the *IER3IP1* p.T79Δ mutant and additional data of surface and total proteome analysis**

**(A)** Control and p.T79Δ mutant-re-expressing cells were treated with 0.66 µM MG132, 33.33 µg/ml chloroquine (CQ) or control DMSO for 24 h, then lysed and analyzed by WB with the indicated antibodies (n = 3 independent experiments). **(B)** mRNA was isolated, and cDNAs were synthesized and analyzed by PCR. **(C)** Control HeLa cells were transfected with mRFP-KDEL, fixed, and labeled with anti-IER3IP1 (green) and Hoechst (blue) (n = 3 independent experiments). Arrows indicate ER membranes co-labeled by IER3IP1. **(D)** Control HeLa cells were co-transfected with mCherry-Sec24C and EGFP-ERGIC53, fixed and stained with anti-IER3IP1 (blue), and analyzed by fluorescence microscopy. Arrows indicate membranes co-labeled by the three proteins. **(E)** Model of IER3IP1 localization relative of ER exit sites, as observed in **(D)**. **(C, D)** Scale bars, 10 µm. Shown are single Apotome sections.

**(F, G)** Quantification of total protein levels of endogenous FGFR3 **(F)** and Unc5B-FLAG **(G)**, as shown in Fig. 3H and Fig. 3J, respectively (n = 6 independent experiments; median and IQR values, Kruskal-Wallis with Dunn’s post-hoc test). **(H, I)** Analysis of Unc5B-FLAG glycosylation. **(H)** Indicated cell lines were transfected with Unc5B-FLAG, lysates were collected after 24 h, treated with EndoH or PNGaseF enzymes overnight, and immunoblotted with anti-FLAG and anti-beta-actin as loading control. * shows the mature, upper band of Unc5B-FLAG that is resistant to EndoH but not to PNGAseF, ** shows the immature, lower band, detected at a lower molecular weight following EndoH deglycosylation. **(I)** Quantification of mature/total Unc5B-FLAG (n = 3 independent experiments, mean ± SD, One-way Anova with Dunnett’s post-hoc test).

**(J)** Volcano plot of proteins identified in the whole cell lysates of control and *IER3IP1* KO1 HeLa cells (see Suppl. Table S3). Proteins significantly different in KO1 *versus* control cells are color-coded as shown in the legend: early/late endosome (red) and cell growth (yellow), transport vesicle (blue) and cell junction (green)-associated. Large circles indicate modifications rescued by the re-expression of the WT protein (n = 4 independent experiments). **(K-N)** Pathway enrichment analysis of hits **(K, L)** decreased (red) or **(M, N** increased (blue) in the cell lysates of *IER3IP1* KO1 compared to control cells.

**(O, P)** Quality control of MS analyses; mean ± SD of Pearson correlations of log2 PG. Quantities measured for the **(O)** surface proteome and **(P)** secretome.

**Supplementary Figure S2. *IER3IP1* KO, *IER3IP1* p.L78P and p.T791′ mutants delay Golgi assembly. (A-C)** Cells were treated with DMSO or 5 µg/ml BFA for 1 h, then either fixed, or washed in fresh medium for 2 h following BFA treatment and fixed. Cells were then stained with anti-ST6GAL1 (green), anti-GM130 (red) and Hoechst (blue) and analyzed by fluorescence microscopy. Scale bars, 10 µm. Shown are single Apotome sections. **(D)** The fraction of cells in which the Golgi apparatus re-assembled after the 2 h wash is shown. Each small circle represents a different image and filled circles the mean values of independent experiments, shown in different colors (n = 3 independent experiments, n _Control_ = 593, n _KO1_ = 1004, n _KO1+WT_ = 906, n _KO1+p.L78P_ = 2390, n _KO1+p.V21G_ = 744, n _KO1+p.T79Δ_ = 1036 cells). **(E)** Co-localization between GM130 and ST6GAL1 was evaluated by measuring the object-based Pearson’s correlation coefficient using Cell Profiler. Data are shown for each object (small circles), and mean values for independent experiments (color-coded filled circles) (n = 3 independent experiments, n _Control_ = 845 cells, 6211 objects, n _KO1_ = 682 cells, 9981 objects, n _KO1+WT_ = 682 cells, 4362 objects, n _KO1+p.L78P_ = 888 cells, 13896 objects, n _KO1+p.V21G_ = 485 cells, 4735 objects, n _KO1+p.T79Δ_ = 535 cells, 11127 objects). **(D, E)** mean ± SD, One-way Anova with Dunnett’s post-hoc test.

**Supplementary Figure S3. The absence of functional IER3IP1 increased the area of ER sheets but did not induce significant ER-stress. (A)** Cells were transfected with EGFP-p180 as a marker for ER sheets and analyzed after 24 h using live fluorescence microscopy. Images are shown as 3D Max Projections of Z-sections (0.26 µm per Z-section) were analyzed using Fiji. Scale bars, 10 µm. **(B)** The ratios between the area covered by EGFP-p180 and the total cell area are shown. Each circle represents a cell, color-coded mean values for each independent are shown as filled circles (n = 3 independent experiments, mean ± SD, One-way Anova with Dunnett’s post-hoc test; n _Control_ = 97, n _KO1_ = 129, n _KO1+WT_ = 104, n _KO1+p.L78P_ = 131 cells).

**(C-F)** Cells overexpressing FGFR5-V5 or control V5 **(C, D)**, or treated with 1 µM thapsigargin or control DMSO for 24 h **(E, F**) were analyzed by immunoblotting (n = 3 independent experiments, mean ± SD, two-tailed Welch’s t-test). **(G-I)** Cells treated with thapsigargin or control DMSO for 4 h were analyzed by immunoblotting with the indicated antibodies. **(G)** n = 3, **(H-I)** n = 5 independent experiments, mean ± SD, One-way Anova with Dunnett’s post-hoc test, and two-tailed Welch’s t-test.

**Supplementary Figure S4. *IER3IP1* KO1 and re-expressed *IER3IP1* p.L78P increase the number of active lysosomes. (A-D)** Cells were **(A)** incubated with Magic Red cathepsin L and Hoechst 33342 dye for 60 min and directly imaged, or **(B)** fixed and labeled with anti-LAMP1 and Hoechst 33342, then analyzed by high-content confocal fluorescence microscopy. Single sections are shown in **(A)**, max projections of Z-stacks are shown in **(B)**. Scale bars, 10 µm. **(C, D)** The number of objects per image was calculated using Cell Profiler and normalized to the number of nuclei. Color-coded small circles represent mean values per well, and filled, large circles depict the means of each independent experiment (n = 4, mean ± SD; One-way Anova with Dunnett’s post-hoc test. **(C)** n _Control_ = 34 363 cells, 19 images; n _KO1_ = 29 720 cells, 20 images; n _KO1+WT_ = 31 535 cells, 20 images; n _KO1+p.L78P_ = 28 188 cells, 19 images; **(D)** n _Control_ = 37 847 cells, 20 images; n _KO1_ = 39 965 cells, 23 images; n _KO1+WT_ = 39 277 cells, 23 images; n _KO1+p.L78P_ = 49 706 cells, 23 images.

**(E-G)** Cells were treated with 500 nM Torin 1 and EBSS starvation medium, or 50 µg/ml chloroquine (CQ) for 5h, then lysed and analyzed by immunoblotting with anti-p62, anti-LC3B and anti-beta-actin as loading control. **(F, G)** The protein amounts of LC3II **(F)** and p62 **(G)** normalized to beta-actin are shown as fold change over the respective non-treated cell line. **(F)** n = 3, **(G)** n = 4, **(F, G)** mean ± SD, two-tailed Mann-Whitney test.

**Supplementary Figure S5. Analyses of the *IER3IP1* p.L78P cell secretome, and of KDELR1_EGFP and KDELR3-EGFP localization. (A)** Volcano plot of proteins secreted by cell lines expressing *IER3IP1* p.L78P or WT (Suppl. Table S5). Proteins whose secretion was modified in p.L78P *vs.* WT cells are shown in red (increased), or blue (decreased). Gene names are shown for some of the proteins mentioned in the text (n _WT_ = 3, n _p.L78P_ = 4). **(B-E)** GSEA pathway enrichment analysis (reactome, GO cellular component/GOCC) of proteins whose secretion was either decreased (red) or increased (blue) in *IER3IP1* p.L78P-compared to *IER3IP1* WT expressing cells.

**(F-I)** Localization of KDELR1-EGFP and KDELR3-EGFP. Cells were transfected with **(F)** KDELR1-EGFP or **(G)** KDELR3-EGFP, fixed, co-labeled with anti-GM130 (red) and Hoechst (blue) and analyzed by fluorescence microscopy. Scale bars, 10 µm. Shown are single Apotome sections. (**H, I)** Ratios of KDELR1-EGFP **(H)** or KDELR3-EGFP **(I)** fluorescence in the Golgi region and the entire cell were calculated using Fiji. Color-coded mean values of independent experiments (large, filled circles,) and values calculated for individual cells (small circles) are shown (n _KDELR1-EGFP_ = 4 and n _KDELR1-EGFP_ = 6 independent experiments; mean ± SD, One-way Anova with Dunnett’s post-hoc test; KDELR1-EGFP: n _Control_ = 216, n _KO1_ = 238, n _KO2_ = 166, n _KO1+WT_ = 135, n _KO1+p.L78P_ = 156 cells; KDELR3-EGFP: n _Control_ = 434, n _KO1_ = 575, n _KO2_ = 289, n _KO1+WT_ = 172, n _KO1+p.L78P_ = 294 cells).

## Supplementary Tables

**Supplementary Table S1. Specific enrichment of biotinylated surface proteins among surface proteins.**

**Sheet 1.** List of surface proteins enriched in biotinylated vs non-biotinylated control cells (highlighted in grey). Proteins unchanged between the two conditions not highlighted.

**Sheet 2.** List of surface proteins enriched in biotinylated control cells identified with the keyword “Glycoproteins” (www.uniprot.org).

**Sheet 3.** List of surface proteins enriched in biotinylated control cells identified with the keyword “Transmembrane” (www.uniprot.org).

**Sheet 4.** List of surface proteins enriched in biotinylated control cells identified with the keywords “plasma membrane”, “cell surface”, “extracellular space”, “extracellular matrix”, “extracellular exosome (www.uniprot.org) and confirmed using https://compartments.jensenlab.org/Search.

**Sheet 5.** List of surface proteins enriched in biotinylated control cells identified with the keyword “Signal peptide” (www.uniprot.org).

**Supplementary Table S2. MS analysis of proteins differentially expressed on cell surface in *IER3IP1* KO1 *versus* control cells.**

**Sheet 1.** List of proteins whose surface expression is modified in *IER3IP1* KO1 *vs.* control cells (volcano plot shown in Fig. 3A). Proteins whose surface levels are increased (blue), reduced (pink) or are not changed (no highlighting) between the two conditions are shown.

**Sheet 2.** List of proteins whose surface expression is modified in *IER3IP1* KO1 *vs.* control cells, rescued by re-expression of *IER3IP1* WT (i.e., expression is not significantly changed in *IER3IP1* WT *vs.* control cells) (Fig. 3A, C).

**Sheet 3.** List of proteins whose surface expression is modified in *IER3IP1* WT *vs.* control cells. Ratios between *IER3IP1* WT and control cells are shown (increased, blue; reduced, pink; unchanged, no highlighting).

**Sheet 4.** GSEA GO Cell Component analysis of proteins whose surface levels are decreased in *IER3IP1* KO1 *vs.* control cells (Fig. 3D).

**Sheet 5.** GSEA GO Cell Component analysis of proteins whose surface levels are increased in *IER3IP1* KO1 *vs.* control cells (Fig. 3F).

**Sheet 6.** GSEA Reactome analysis of proteins whose surface levels are decreased in *IER3IP1* KO1 *vs.* control cells (Fig. 3E).

**Sheet 7.** GSEA Reactome analysis of proteins whose surface levels are increased in *IER3IP1* KO1 *vs.* control cells (Fig. 3G).

**Supplementary Table S3. Total proteome analysis in *IER3IP1* KO1 *versus* control cells.**

**Sheet 1.** List of proteins whose total cellular expression is modified in *IER3IP1* KO1 *vs.* control cells (Suppl. Fig. S1J). Ratios between *IER3IP1* KO1 and control cells are shown (increased, blue; reduced, pink; unchanged, not highlighted).

**Sheet 2.** GSEA GO Cell Component analysis of proteins whose total levels are decreased in *IER3IP1* KO1 *vs.* control cells (Suppl. Fig. S1J, K).

**Sheet 3.** GSEA GO Biological Process analysis of proteins whose total levels are decreased in *IER3IP1* KO1 *vs.* control cells (Suppl. Fig. S1J, L).

**Sheet 4.** GSEA GO Cell Component analysis of proteins whose total levels are increased in *IER3IP1* KO1 *vs.* control cells (Suppl. Fig. S1J, M).

**Sheet 5.** GSEA Reactome analysis of proteins whose total levels are increased in *IER3IP1* KO1 *vs.* control cells (Suppl. Fig. S1J, N).

**Supplementary Table S4. Specific enrichment of biotinylated secreted proteins.**

**Sheet 1.** List of secreted proteins enriched in biotinylated vs non-biotinylated control cells. Proteins significantly modified are highlighted in blue, and those unchanged are not highlighted.

**Sheet 2.** List of N-Glycosylated proteins identified with the keyword “Glycoproteins” (www.uniprot.org) among the enriched biotinylated proteins.

**Sheet 3.** List of proteins identified among the enriched biotinylated proteins with the keywords “plasma membrane”, “cell surface”, “extracellular space”, “extracellular matrix”, “extracellular exosome (www.uniprot.org) and confirmed using https://compartments.jensenlab.org/Search.

**Sheet 4.** List of proteins identified among the enriched biotinylated proteins with the keyword “Secreted” (www.uniprot.org).

**Sheet 5.** List of proteins identified among the enriched biotinylated proteins with the keyword “Transmembrane” (www.uniprot.org).

**Sheet 6.** List of proteins identified among the enriched biotinylated proteins with the keyword “Signal peptide” (www.uniprot.org).

**Supplementary Table S5. MS analysis of differentially secreted proteins in *IER3IP1* KO1 *versus* control cells.**

**Sheet 1.** List of proteins whose secretion is modified in *IER3IP1* KO1 *vs.* control cells (Fig. 4A). Ratios between *IER3IP1* KO1 and control cells are shown (increased, blue; reduced, pink; unchanged, not highlighted).

**Sheet 2.** List of proteins whose secretion is modified in *IER3IP1* KO1 *vs.* control cells, and rescued by re-expression of *IER3IP1* WT (i.e., expression is not significantly changed in *IER3IP1* WT *vs.* control cells) (Fig. 4A, C). Proteins partially rescued are highlighted in light orange.

**Sheet 3.** List of proteins whose secretion is modified in *IER3IP1* WT *vs.* control cells. Ratios between *IER3IP1* WT and control cells are shown in blue (increased) pink (decreased) or were not highlighted (unchanged).

**Sheet 4.** List of proteins whose secretion is modified in *IER3IP1* p.L78P *vs.* WT cells (Suppl. Fig. S5A). Proteins are highlighted in blue (increased), pink (decreased) or not highlighted (unchanged). Proteins that are differentially secreted in *IER3IP1* KO1/Control and *IER3IP1* p.L78P/*IER3IP1* WT are shown in bold.

**Sheet 5.** GSEA Reactome analysis of proteins whose secretion is decreased in *IER3IP1* KO1 *vs.* control cells (Fig. 4A, D).

**Sheet 6.** GSEA GO Cell Component analysis of proteins whose secretion is decreased in *IER3IP1* KO1 *vs.* control cells (Fig. 4A, E).

**Sheet 7.** GSEA Reactome analysis of proteins whose secretion is increased in *IER3IP1* KO1 *vs.* control cells (Fig. 4A, F).

**Sheet 8.** GSEA GO Cell Component analysis of proteins whose secretion is increased in *IER3IP1* KO1 *vs.* control cells (Fig. 4A, G).

**Sheet 9.** GSEA Reactome analysis of proteins whose secretion is decreased in *IER3IP1* p.L78P *vs.* WT cells (Suppl. Fig. S5A, B).

**Sheet 10.** GSEA GO Cell Component analysis of proteins whose secretion is decreased in *IER3IP1* p.L78P *vs.* WT cells (Suppl. Fig. S5A, C).

**Sheet 11.** GSEA Reactome analysis of proteins whose secretion is increased in *IER3IP1* p.L78P *vs.* WT (Suppl. Fig. S5A, D).

**Sheet 12.** GSEA GO Cell Component analysis of proteins whose secretion is increased in *IER3IP1* p.L78P *vs.* WT cells (Suppl. Fig. S5A, E).

**Supplementary Table S6. ER retention motifs.** List of proteins identified with the motifs “Prevents secretion from ER” or “Non-canonical ER retention motif” (www.uniprot.org) among the proteins whose secretion is higher in *IER3IP1* KO1 cells compared to control.

**Supplementary Table S7. Baso-lateral and apical proteins** whose surface expression or secretion was modified in *IER3IP1* KO1 cells compared to controls. Polarized protein distribution was analyzed with http://polarprotdb.ttk.hu/search^86^.

