## Supplementary figures and images for "IER3IP1-mutations cause microcephaly by selective inhibition of ER-Golgi transport"

### Supplemental figures S1-S5

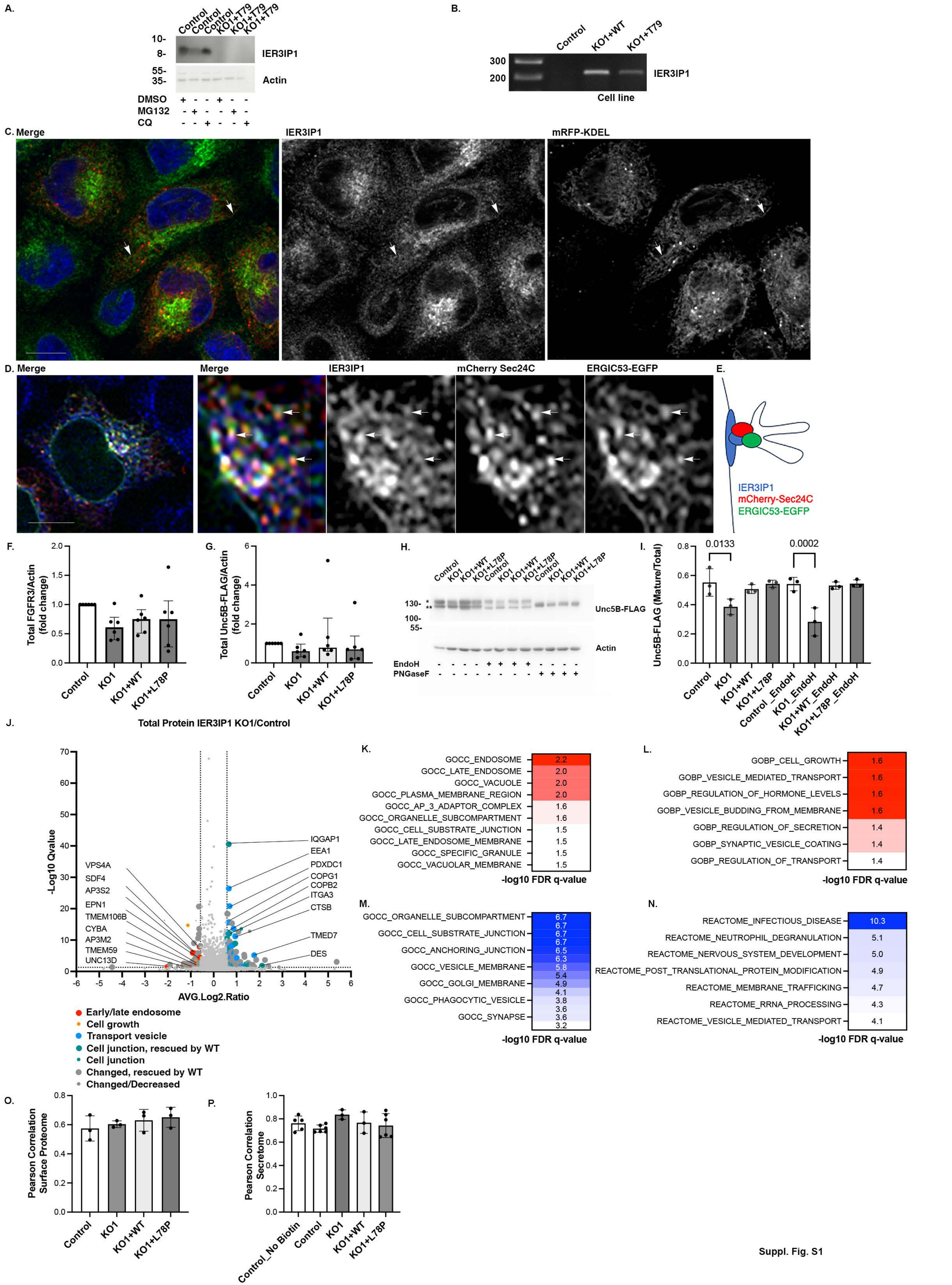

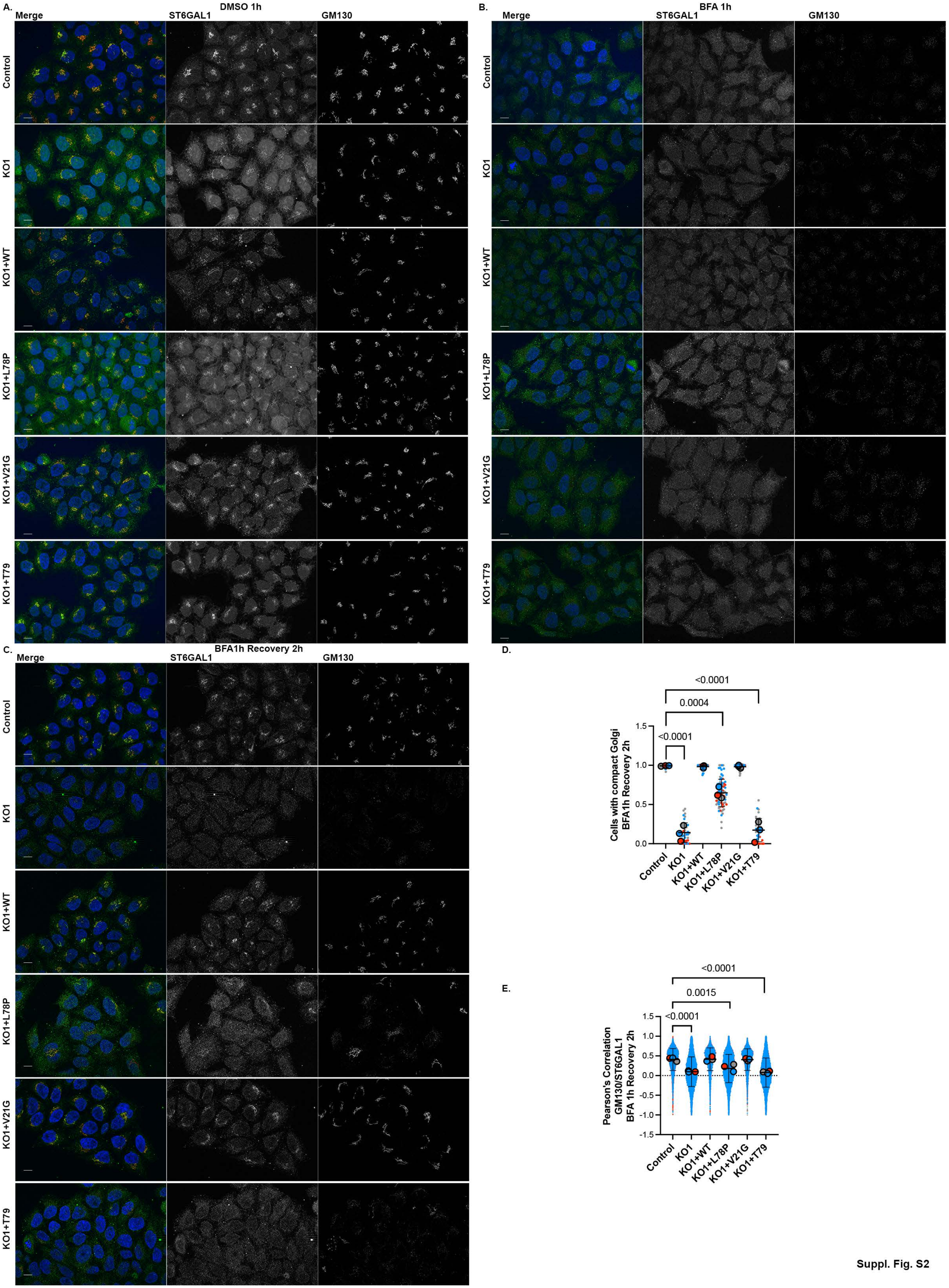

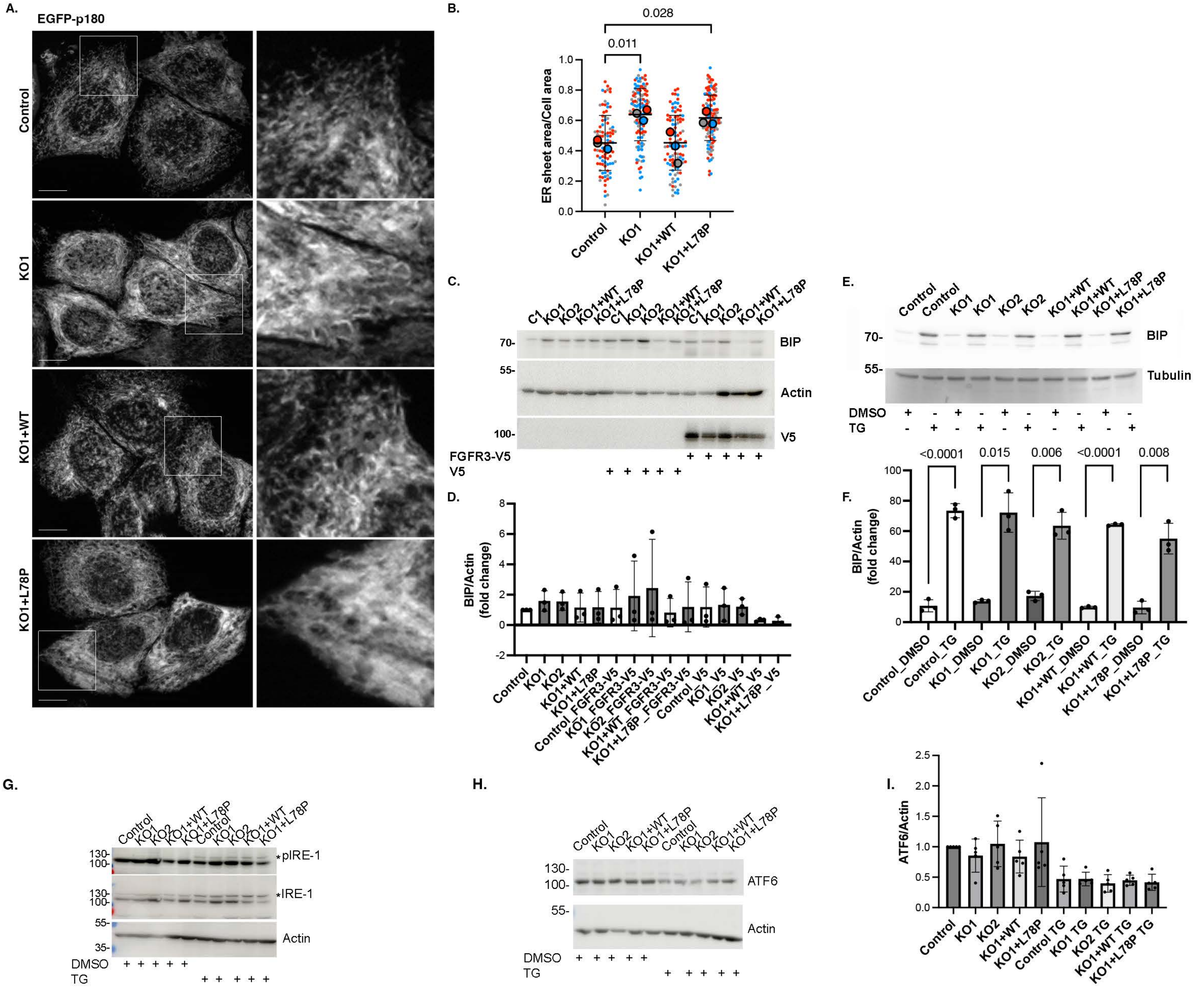

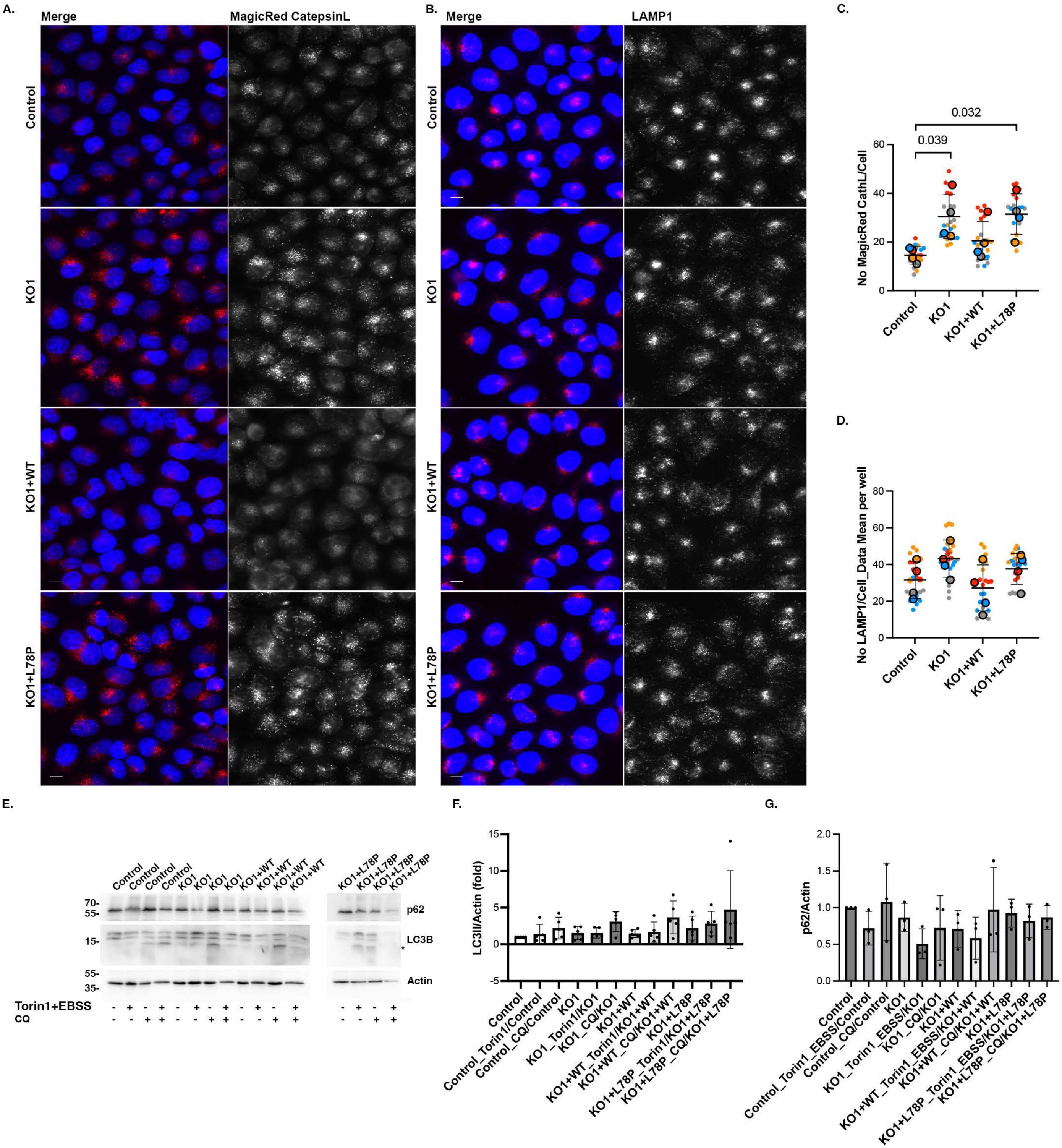

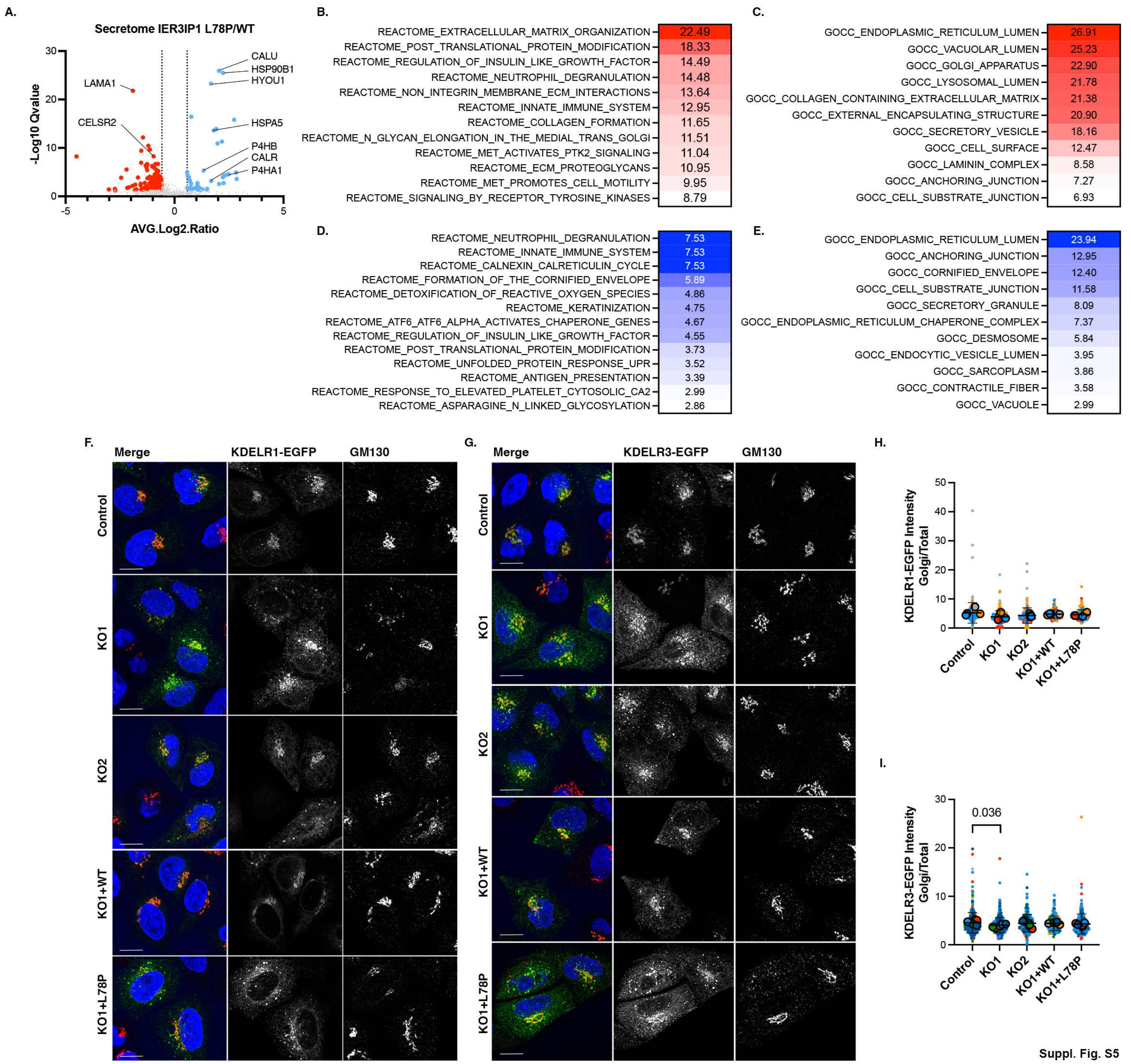
